# SMCHD1 maintains heterochromatin and genome compartments in human myoblasts

**DOI:** 10.1101/2024.07.07.602392

**Authors:** Zhijun Huang, Wei Cui, Ishara Ratnayake, Rabi Tawil, Gerd P. Pfeifer

## Abstract

Mammalian genomes are subdivided into euchromatic A compartments that contain mostly active chromatin, and inactive, heterochromatic B compartments. However, it is unknown how A and B genome compartments are established and maintained. Here we studied SMCHD1, an SMC-like protein in human male myoblasts. SMCHD1 colocalizes with Lamin B1 and the heterochromatin mark H3K9me3. Loss of SMCHD1 leads to extensive heterochromatin depletion at the nuclear lamina and acquisition of active chromatin states along all chromosomes. In absence of SMCHD1, long range intra-chromosomal and inter-chromosomal contacts between B compartments are lost while many new TADs and loops are formed. Inactivation of SMCHD1 promotes numerous B to A compartment transitions accompanied by activation of silenced genes. SMCHD1 functions as an anchor for heterochromatin domains ensuring that these domains are inaccessible to epigenome modification enzymes that typically operate in active chromatin. Therefore, A compartments are formed by default when not prevented by SMCHD1.

## INTRODUCTION

The genome of eukaryotes consists of segments of euchromatin and heterochromatin. Euchromatin contains transcriptionally active genomic regions whereas heterochromatin is largely depleted of genes and contains transcriptionally silent sequences and repetitive DNA ^1–6^. Histone H3 lysine 9 (H3K9) methylation is a hallmark of heterochromatin ^3^. Almost half of the human genome consists of heterochromatin, but the structural and functional role of heterochromatin is only partially understood ^7–9^. At the three-dimensional level, mammalian chromosomes are roughly subdivided into A compartments that contain mostly active chromatin and B compartments that consist chiefly of heterochromatin ^10–13^. However, it is not clear how these compartments are established and maintained.

Structural maintenance of chromosomes flexible hinge domain containing 1 (SMCHD1) was initially identified in a genetic screen as a repressor of mouse metastable epialleles ^14^. This protein is involved in X chromosome inactivation by facilitating the hypermethylation of CpG islands and by other chromatin-based repression mechanisms ^14–16^. SMCHD1 is mutated in two unrelated human genetic diseases, facioscapulohumeral muscular dystrophy type 2 (FSHD2) ^17,18^ and Bosma arrhinia microphtalmia syndrome (BAMS) ^19,20^, a rare developmental defect. The FSHD2 alterations of SMCHD1 are generally considered loss of function mutations whereas this has been less clear for the BAMS mutations ^16^. Although the function of SMCHD1 in X chromosome inactivation is quite well characterized ^21–27^, we know little about the role of this protein in the control of chromatin structure on autosomes. Previous work has shown that SMCHD1 is as a regulator of *HOX* gene clusters, protocadherin clusters and of imprinted gene expression ^15,28–32^, but its role is thought to be restricted to a very limited number of autosomal gene regions.

However, given the ubiquitous and cell type independent expression of SMCHD1 in male and female cells, we reasoned that SMCHD1 may have roles that could go well beyond regulating the X chromosome and specific gene loci. Because of potential disease relevance, we focused here on human myoblast cells. We characterized the genomic properties of SMCHD1 as a nuclear lamina-associated protein. Using gene inactivation in combination with detailed genome-wide DNA methylation and chromatin structure analysis including HiC mapping, we show that loss of SMCHD1 in these somatic human cells leads to a genome-wide perturbation of heterochromatin structure, whereby many heterochromatic B compartment regions loose contact to each other and undergo transition to euchromatic A compartments accompanied by major changes in 3D chromatin structure and epigenome landscape.

## RESULTS

### SMCHD1 colocalizes with Lamin B1 and the heterochromatin mark H3K9me3

The male immortalized human myoblast cell line LHCN-M2 was grown under standard conditions without differentiation. Using these cells, we performed a comprehensive characterization of SMCHD1 and its relationship to DNA methylation, chromatin modifications, gene expression, and three-dimensional genome structure (Fig. 1A). To map the genomic distribution of SMCHD1, we used Dam-ID, a method in which a small adenine methyltransferase domain is attached to the 3’ end of the SMCHD1 protein ^23^. The methylated adenines, which are established on DNA in the vicinity of the tagged protein, are then mapped using cleavage with the 6-methyladenine-dependent restriction enzyme DpnI and high-throughput sequencing (Fig. 1B). Inspection of the Dam-ID SMCHD1 signals along all chromosomes showed a generally broad distribution of the protein in large megabase-size blocks that were interrupted by areas of much weaker SMCHD1 signal (Fig. 1B; Extended Data Table 1). The Dam-only sequencing controls showed only a low and non-specific signal along the genome and the SMCHD1 signal was normalized relative to the Dam only signal (Fig. 1B). Known SMCHD1-bound genes such as the protocadherin clusters and imprinted genes showed smaller SMCHD1 peaks. The large SMCHD1 blocks tended to be localized in gene-poor areas of the chromosomes. Because gene-depleted areas of the genome are known to be associated with the nuclear lamina and with heterochromatin and transcriptional repression ^9,33–35^, we mapped in parallel the distribution of Lamin B1 by Dam-ID sequencing and of the characteristic heterochromatin mark histone H3 lysine 9 trimethylation (H3K9me3) by standard ChIP sequencing (Fig. 1B, Fig. 1C-E; Extended Data Fig. 1). SMCHD1, Lamin B1, and H3K9me3 were extensively colocalized (Fig. 1B-H; Extended Data Fig. 1A, 2C). Heat maps show the clear colocalization of the three mapped features over lamina-associated domain (LAD) regions (Fig. 1C-E). Venn diagrams (Fig. 1F) and scatter plots (Fig. 1H) further reveal the strongly correlated localization of SMCHD1 and Lamin B1. Colocalization was found along all chromosomes (Extended Data Fig. 1A). Only chromosomes 19 and 22, two gene-rich chromosomes, did not contain many SMCHD1 and Lamin B1 blocks (Extended Data Fig. 1A). The distributions of SMCHD1, Lamin B1, and H3K9me3 over LAD borders showed a sharp rise for SMCHD1 and Lamin B1 and a more gradual increase of H3K9me3 from the borders into the LADs themselves (Extended Data Fig. 1B). As expected, the euchromatic (active) marks H3K4me1, H3K4me3, H3K27 acetylation, H3K36me2, H3K36me3, and RNA polymerase II, which we also mapped in the LHCN-M2 cells, were depleted within the LAD regions (Extended Data Fig. 1B). Principal component analysis (PCA) shows the distinct differences between the SMCHD1-bound and non-bound regions based on the mapped histone modification patterns and Lamin B1 binding pattern (Fig. 1G). Extended Data Fig. 1C shows an example of the different chromatin modifications present at SMCHD1-bound and SMCHD1-unbound regions and Extended Data Fig. 1D summarizes the differential distributions of these marks.

**Figure 1:**
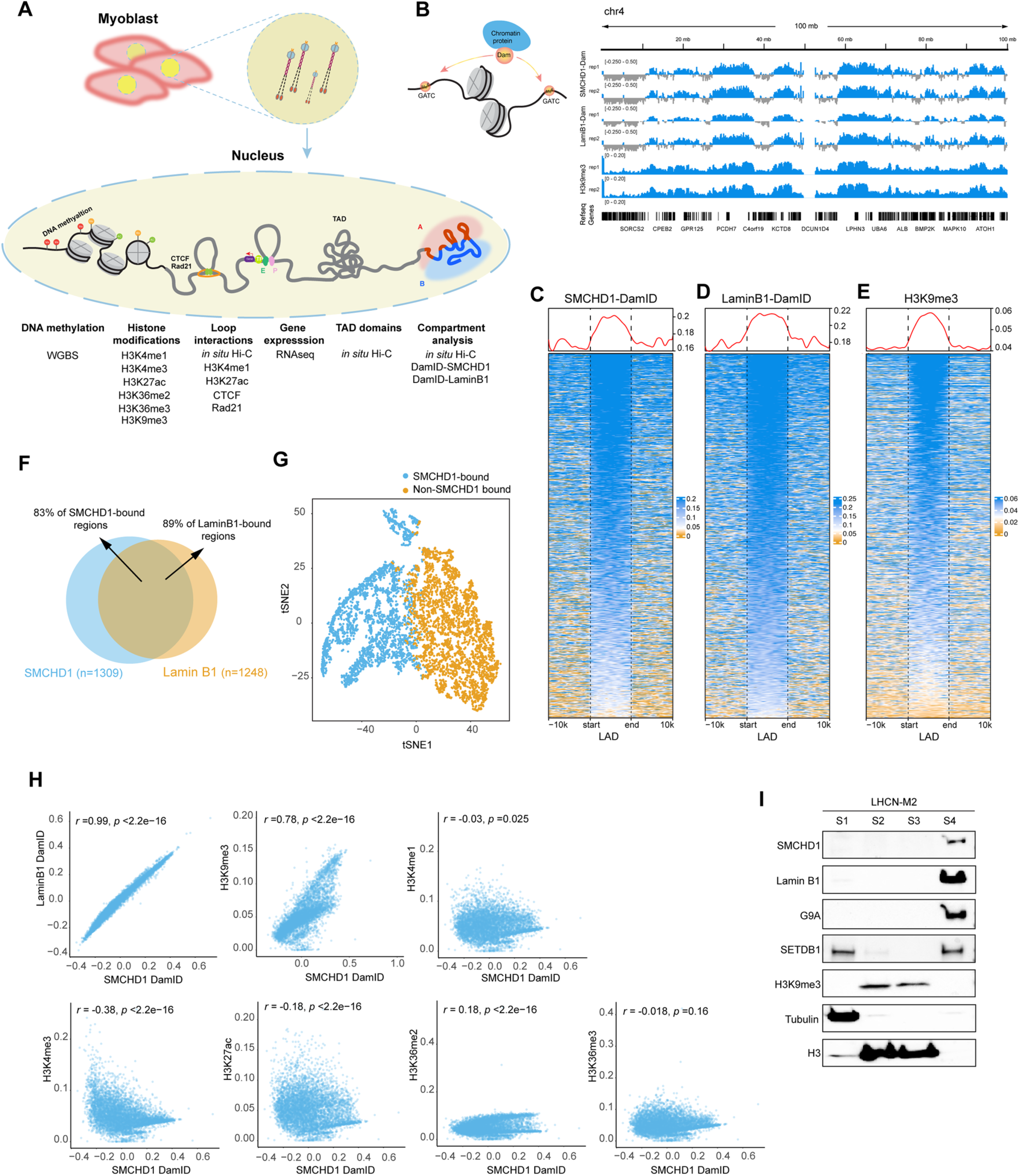
Colocalization of SMCHD1 with Lamin B1 and H3K9me3. **A**. Outline of the experimental approaches used in this study. **B.** Dam-ID sequencing shows the localization of SMCHD1 and Lamin B1 along chromosome 4. Two replicates each are shown. The signals were normalized relative to Dam-only signal. H3K9me3 was mapped by ChIP-sequencing. The density of Refseq genes (bottom) shows that the SMCHD1 blocks preferentially lie within gene-poor regions. **C.** Heat map of SMCHD1-DamID signal over LAD regions. **D.** Heat map of Lamin B1-DamID signal over LAD regions. **E.** Heat map of H3K9me3 ChIP-seq signal over LAD regions. **F.** More than 80% of SMCHD1 and Lamin B1 regions coincide. **G.** tSNE plots of SMCHD1-bound and SMCHD1-unbound bins (500 kb) at the whole genome level. All bins were plotted based on the combinatorial mean of H3K9me3, H3K4me1, H3K4me3, H3K27ac, H3K36me2, H3K36me3 and Lamin B1 levels across all bins in two dimensions, using t-Distributed Stochastic Neighbor Embedding (t-SNE). **H.** Scatter plots show a strong positive correlation between SMCHD1 and Lamin B1 DamID signals and H3K9me3 but marginal or negative correlation with active chromatin marks. **I.** Cell fractionation experiments and Western blots showing the enrichment of SMCHD1, Lamin B1, G9a (EHMT2), SETDB1, H3K9me3, tubulin, and total histone H3 in cytoplasmic (S1), soluble chromatin (S2), tightly chromatin-associated (S3) and insoluble cellular fractions (S4) of LHCN-M2 cells.

To obtain further support for the association of SMCHD1 with the nuclear lamina, we performed cell fractionation experiments. We tested the presence of SMCHD1, Lamin B1, the H3K9 methyltransferases G9A (EHMT2) and SETDB1 and of H3K9me3 in cytoplasmic (S1), soluble chromatin (S2), tightly chromatin-associated (S3), and insoluble (lamina-containing) (S4) cellular fractions by Western blotting (Fig. 1I). While the histones were extracted into the S2 and S3 fractions, SMCHD1, Lamin B1 and the H3K9 methylation enzymes remained in the insoluble S4 fraction consistent with our genomic mapping data (Fig. 1I).

### SMCDH1 maintains heterochromatin architecture at the nuclear lamina

To study the function of SMCHD1 in muscle cells, we used CRISPR/Cas9 technology to inactivate this gene. Figure 2A shows the complete loss of SMCHD1 protein in three independent knockout cell clones relative to three wildtype cell clones, which were subsequently used throughout the study. Extended Data Figure 2A shows the genotypes of the knockout clones. We used transmission electron microscopy (TEM) to monitor structural changes of the nucleus in SMCHD1-deficient cells. In wildtype myoblasts (Fig. 2B), we observed dense clusters of heterochromatin at the nuclear periphery, as expected. However, in cells lacking SMCHD1, a much thinner layer of heterochromatin at the outer boundaries of the nucleus was observed (Fig. 2C). Densitometric quantification of the heterochromatin layer near the nuclear surface shows a significant reduction of staining intensity in the knockout cells (P= 4.288e-06) (Fig. 2D). Next, we analyzed myoblast cells obtained from FSHD2 patients carrying heterozygous loss of function mutations in SMCHD1 (Extended Data Fig. 2). Like in the immortalized myoblasts with inactive SMCHD1, we find a reduction of heterochromatin density at nuclear lamina attached regions in myoblasts from FSHD2 patients (P = 2.184e-10, t-test; Extended Data Fig. 2B-D).

**Figure 2:**
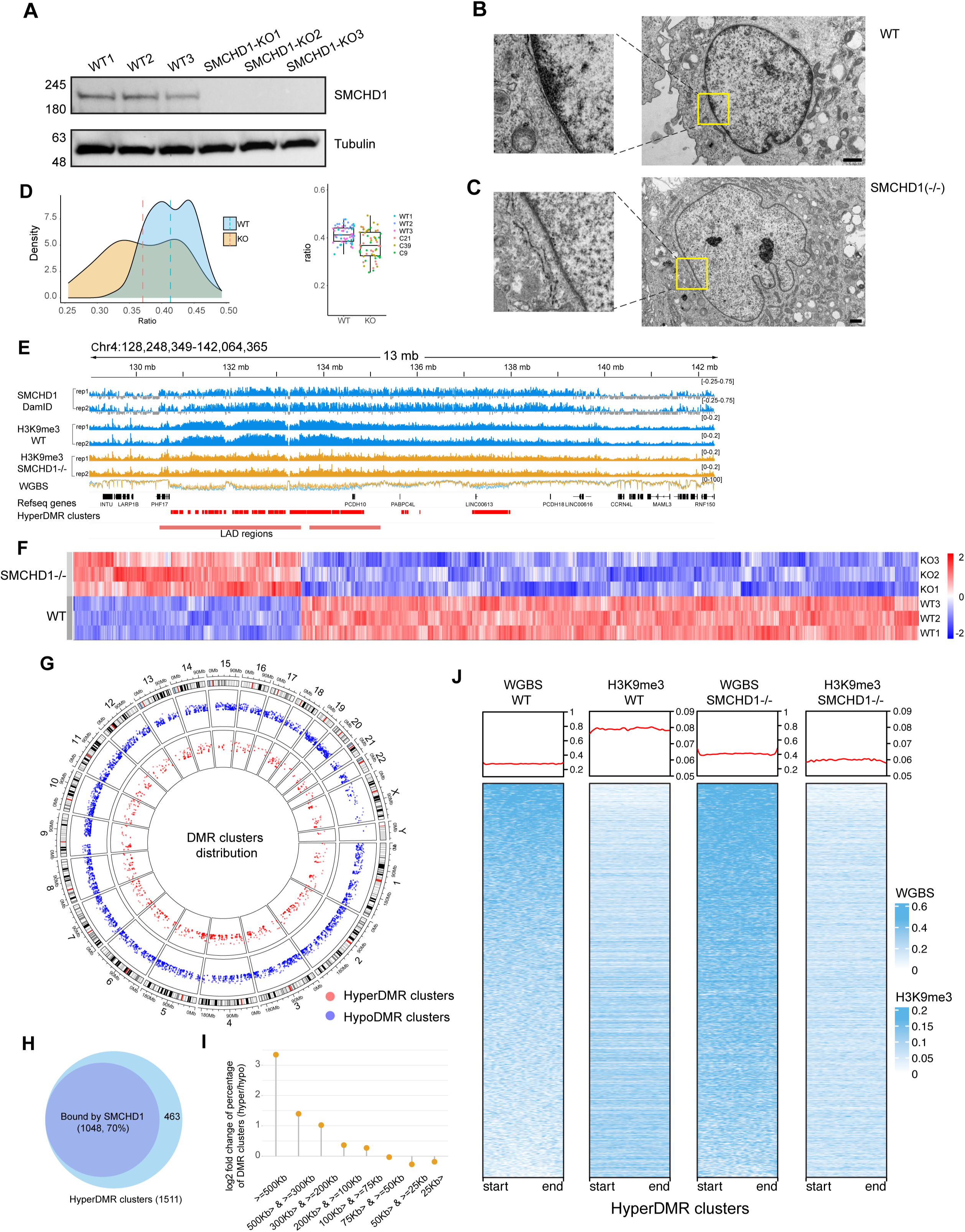
Inactivation of SMCHD1 leads to changes in heterochromatin, H3K9me3 and DNA methylation patterns. **A**. CRISPR/Cas9-mediated inactivation of SMCHD1 in LHCN-M2 myoblasts. Three wildtype and three SMCHD1-deficient clones were analyzed by Western blot. **B.** Transmission electron microscopy (TEM) shows dense heterochromatin staining at the nuclear periphery in wildtype (WT) cells. The magnified squares show the nuclear envelope regions in more detail. **C.** TEM shows much weaker staining of heterochromatin at the nuclear periphery in SMCHD1 knockout cells. **D.** Densitometric quantitation of nuclear lamina-associated heterochromatin regions in WT and SMCHD1 KO cells. Regions under the nuclear envelope were quantitated by densitometry of 59 WT and 58 KO cells. The heterochromatin staining intensities were normalized to the total area of the nucleus. P=4.288e-06, t-test. **E.** Analysis of H3K9me3 by ChIP-seq and of DNA methylation by whole genome bisulfite sequencing (WGBS) in wildtype and SMCHD1-deficient myoblasts. Genome browser views of a region of chr4 are shown. The localization of SMCHD1 is shown in the top two tracks. Refseq genes, hypermethylation DNA clusters, and LAD regions are shown at the bottom. Overlaid tracks for WGBS are shown for wildtype (blue) and SMCHD1^−/−^ cells (orange). **F.** Heat map of differentially methylated regions (DMRs) between SMCHD1 knockout (KO) and wildtype (WT) cells. **G.** Circos plot showing hypomethylation DMR (hypoDMR, blue) and hypermethylation DMR (hyperDMR, red) clusters along all chromosomes. **H.** Overlap between SMCHD1-bound regions and DNA hypermethylation clusters. **I.** Hypermethylation DMR clusters have greater length (in kb) than hypomethylation DMR clusters. **J.** Heatmaps for DNA methylation levels (WGBS) and H3K9me3 signals over all hypermethylation DMR cluster regions in WT and SMCHD1^−/−^ cells. Density plots are shown near the top.

SMCHD1 has previously been linked to H3K9 trimethylation ^36,37^ suggesting a connection between SMCHD1, likely via the bridging proteins LRIF1/HBIX1 and HP1, and the constitutive heterochromatin histone mark H3K9me3. The colocalization of SMCHD1 and H3K9me3 we observe in human muscle cells (Fig. 1B-H; Extended Data Fig. 1) supports a tight connection between SMCHD1 and constitutive heterochromatin. Inspection of genome browser data shows that many regions with megabase-size SMCHD1 blocks in wildtype cells undergo a loss of H3K9me3 in the SMCHD1 knockout cells (Fig. 2E; Extended Data Fig. 3A) supporting a role of SMCHD1 in heterochromatin integrity. Overall, we calculated that about 50% of all broad SMCHD1-associated regions as determined by Dam-ID sequencing show a 2-fold or more diminished signal of H3K9me3 in the knockouts.

### Re-organization of DNA methylation in SMCHD1-deficient muscle cells

Since SMCHD1 has previously been implicated in the control of DNA methylation ^14,28,30,38^, we performed whole genome bisulfite sequencing of the three wildtype myoblast cell clones and three SMCHD1 knockout clones (Fig. 2E-J; Extended Data Fig. 3A). In total, there were 115,400 hypomethylated differentially methylated regions (DMRs) and 48,500 hypermethylated DMRs in the SMCHD1^−/−^ cells. These DMRs occurred along all chromosomes (Fig. 2G). Upon closer inspection, we found that the hypomethylation DMRs were shorter and more scattered along the genome than the hypermethylation DMRs, which tended to occur in tightly linked long clusters (Fig. 2E, 2I; Extended Data Fig. 3A). The overall preponderance of hypomethylation DMRs over hypermethylation DMRs in terms of numbers is consistent with a role of SMCHD1 as a negative regulator of 5-methylcytosine oxidase enzymes, which catalyze DNA demethylation. These enzymes should be more active in the absence of SMCHD1 as previously reported ^30^.

The hypermethylated regions displayed very unusual and interesting features. First, most hyper-DMRs occurred in dense clusters (n=1,511; Extended Data Table 2) and as such covered larger genomic regions of several megabases. DMR clusters were defined as having five or more DMRs within 10 kb, and the DMR clusters were joined when the maximum distance between the DMR clusters was less than 10 kb. Examples are shown in Figure 2E, Extended Data Fig. 3A, and in subsequent Figures. The hypermethylated DMR clusters were longer (average length 90 kb) than the hypomethylated DMR clusters (average length 59 kb) (Fig. 2I). Second, the hypermethylation DMR clusters (70%) were preferentially associated with SMCHD1-bound regions (Fig. 2H). Third, hypermethylation DMR regions show a concomitant loss of H3K9me3 (Fig. 2E, 2J; Extended Data Fig. 3A). The latter inverse correlation between DNA methylation and the heterochromatin modification H3K9me3 is highly unusual because the two inactive gene marks usually coexist, for example for repression of repetitive DNA sequences ^8,39^. In our system, DNA hypermethylation occurred along LAD regions, bound by SMCHD1. These regions generally tend to be only partially methylated in most cell types and have been referred to as partially methylated domains (PMDs) ^40^. In essence, loss of SMCHD1 converts these partially methylated domains into a state of higher methylation, with concomitant loss of H3K9me3.

### Acquisition of active chromatin marks in SMCHD1 bound regions after loss of SMCHD1

We then mapped several active chromatin marks in wildtype and SMCHD1-depleted cells. The major differences were again found in SMCHD1-bound regions, most notably in those that acquired long stretches of DNA hypermethylation (Extended Data Fig. 3). Regions that acquired clustered DNA hypermethylation and lost H3K9me3 simultaneously gained extensive signals for H3K36me2, H3K36me3, numerous peaks of H3K27 acetylation, H3K4me1 and H3K4me3 (Extended Data Fig. 3A). This observation was made genome-wide for hypermethylation DMR clusters (Extended Data Fig. 3B, 3C). PCA separates the hypermethylated DMR clusters between wildtype cells and SMCHD1-deficient cells when all mapped histone modifications are considered (Extended Data Fig. 3D). For the hypomethylation DMR clusters, the reverse phenomenon was observed, i.e., loss of active marks, but this occurred to a much lesser extent (Extended Data Fig. 3C). These regions showed a limited gain of H3K9me3 (Extended Data Fig. 3C). Hypermethylation DMR clusters in SMCHD1-bound regions with a concomitant loss of H3K9me3 occurred along all chromosomes (Extended Data Fig. 3E-F).

We also mapped the structural proteins CTCF and the cohesin subunit RAD21 in wildtype and SMCHD1 knockout cells. Also here, within the SMCHD1 bound regions and hypermethylation DMR clusters, we observed a strong gain of many coinciding CTCF and RAD21 peaks (Extended Data Fig. 3A-C). Examples are shown in the bottom panels of Extended Data Fig. 3A, and genome-wide analysis is presented in Extended Data Fig. 3B and 3C.

### Control of genome compartmentalization by SMCHD1

The changes in CTCF and cohesin (RAD21 subunit) localization suggested that the three-dimensional genome architecture may undergo substantial changes when SMCHD1 is inactivated. To obtain a detailed picture of 3D genome structure in wildtype and SMCHD1-depleted muscle cells, we used the HiC 3.0 technique ^41,42^ on two biological replicates each of wildtype and SMCHD1 knockout LHCN-M2 myoblasts. We obtained between 2.1 and 2.5 billion aligned contact reads for each sample providing detailed resolution down to a level of a few kilobases (Extended Data Fig. 4A). We pooled the data from the biological replicates, which were well correlated (Extended Data Fig. 4B), to achieve high read density. Figure 3A shows contact maps along the entire chromosome 4 as an example. We observed the loss of many long-range (5 to 10 Mb distance) contacts in the SMCHD1^−/−^ cells (Fig. 3A, 3B). However, when examining the chromosomes at higher magnification, we reveal a gain of shorter-range contacts within windows of single to a few megabases (Fig. 3B). As shown for the entire genome (Fig. 3C), or specifically for chromosome 2 as an example (Fig. 3D), the average probability to gain contacts in the SMCHD1 knockout relative to wildtype cells is increased for contact ranges from 0 to approximately 2 Mb and then declines for the longer-range contacts.

**Figure 3:**
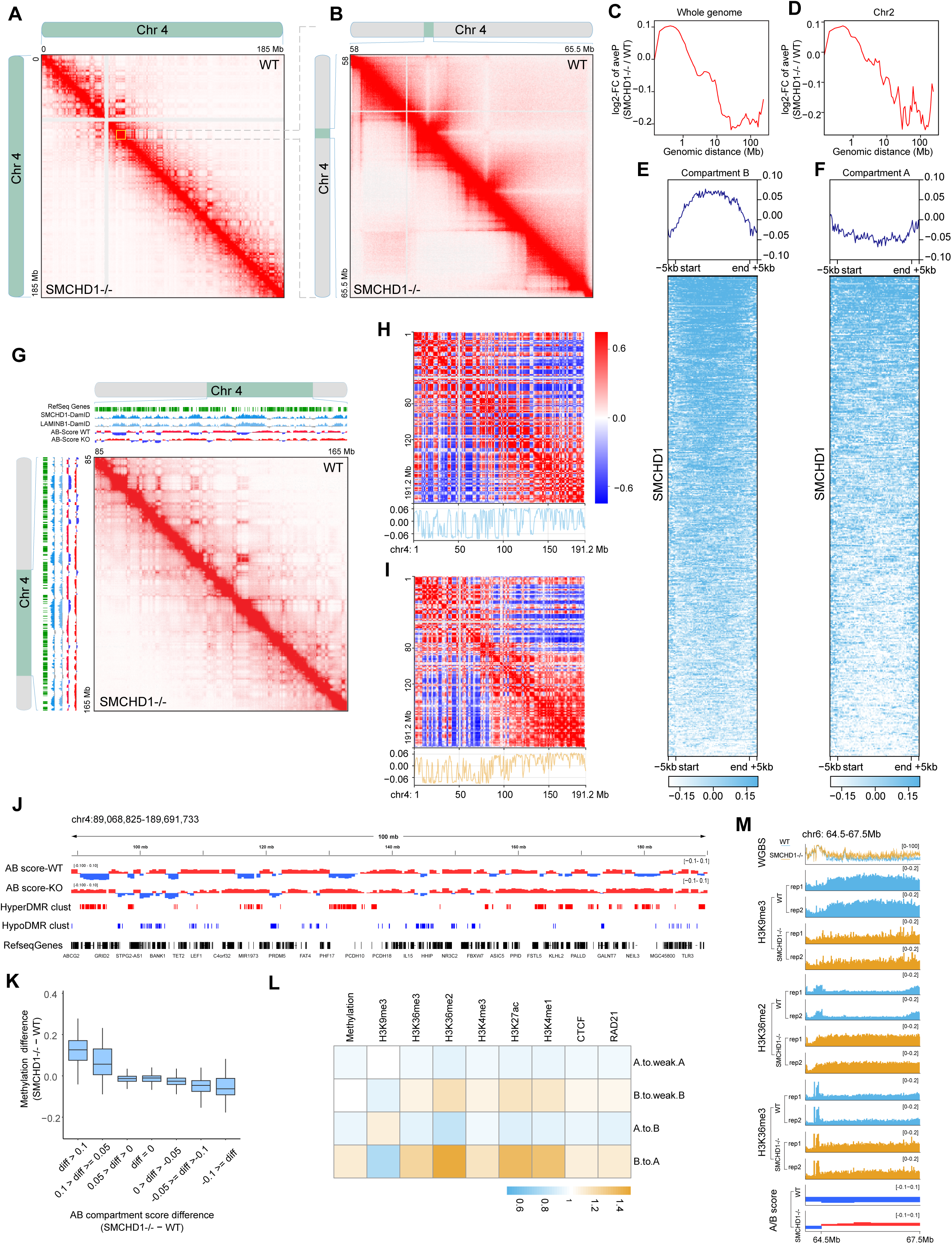
SMCHD1 controls genome compartmentalization. **A**. In situ Hi-C contact maps showing contacts along the entire chromosome 4 in wildtype (WT) and SMCHD1^−/−^ myoblasts. Many long-range contacts are diminished or lost, along with gain of many short-range contacts in the SMCHD1-deficient cells. **B.** Enlargement of an area on chromosome 4 from panel A. This region shows gain of numerous short-range contacts in the SMCHD1^−/−^ cells. **C.** Log-fold change of relative contact probabilities between SMCHD1^−/−^ and wildtype cells depending on genomic distance for the whole genome. **D.** Log-fold change of relative contact probabilities between SMCHD1^−/−^ and wildtype cells depending on genomic distance for chromosome 2. **E.** Distribution of SMCHD1 Dam-ID signal over B compartments derived from HiC analysis. **F.** Distribution of SMCHD1 Dam-ID signal over A compartments derived from HiC analysis. **G.** HiC contacts and Hi-C eigenvector (PC1) analysis in WT and SMCHD1^−/−^ cells across an 80 Mb region of chromosome 4. The SMCHD1 and Lamin B1 signals are shown along with B compartments (blue) and A compartments (red). B-to-A transitions can be seen in the knockout cells compared to wildtype. **H.** Heat maps showing the PC1 for chromosome 4 at 500-kb resolution in wildtype cells. Bottom: Line plot showing corresponding PC1 for chromosome 4. **I.** Heat maps showing the PC1 for chromosome 4 at 500-kb resolution in SMCHD1^−/−^ cells. Bottom: Line plot showing corresponding PC1 for chromosome 4. **J.** Detailed display of A and B compartments in WT and SMCHD1^−/−^ cells along with their positional relationship to hyperDMR clusters, hypoDMR clusters and Refseq genes. **K.** Boxplot showing association between compartment score (Hi-C Eigenvector) change and DNA methylation change. Positive values of AB compartment score difference indicate chromatin activation in association with DNA hypermethylation, negative values of AB compartment score difference indicate chromatin repression in association with DNA hypomethylation. **L.** Heatmap showing changes of chromatin marks, DNA methylation, CTCF and RAD21 in relation to compartment transitions in SMCHD1^−/−^ cells. **M.** Plots showing changes in DNA methylation (WGBS) and chromatin marks over a B-to-A compartment transition for a representative region on chromosome 6.

We next assigned chromatin compartments (A versus B) using Eigenvector principal component analysis ^41^. Most B (inactive chromatin) but not A (active) compartments were associated with SMCHD1 binding (Fig. 3E, 3F). We observed that loss of SMCHD1 promoted many B-to-A compartment transitions (Fig. 3G-J; shift of blue to red color of the compartment scores). These transitions occurred specifically in SMCHD1-associated regions (Fig. 3G). Furthermore, B to A compartment transitions were confirmed when comparing the heatmaps of PC1 over the whole chromosome in WT and SMCHD1^−/−^ cells (Fig. 3H, 3I).

Almost half (n=317) of all B compartments transitioned to A compartments whereas a lower number of A compartments changed into B (Extended Data Fig. 5A) but over much smaller regions (Fig. 3G, 3J; Extended Data Fig. 5B). Of the remaining “non-transitioning” B compartments (N=328), many of them (n=146) became converted into weak B compartments (Extended Data Fig. 5A). This means that over 70% of all B compartments are converting to weak B or A compartments. Most B-to-A transitioning regions contained DNA hypermethylation clusters (Fig. 3J; Extended Data Fig. 5B). The greatest DNA methylation differences were observed at the highest compartment score differences, reflecting B-to-A transitions (Fig. 3K). Globally, B-to-A compartment transitions were associated with a loss of H3K9me3 and with gains of DNA methylation, H3K36me2, H3K36me3, H3K4me1, H3K4me3, H3K27Ac, CTCF and RAD21 (cohesin) (Fig. 3L; Extended Data Fig. 5C). Figure 3M shows an example from chromosome 6, where a B-to-A compartment transition is accompanied by changes in several of these chromatin marks.

### Loss of intrachromosomal and interchromosomal inter-B compartment contacts

Importantly, there were many long-range contacts between different SMCHD1-bound B compartments (B-to-B contacts) along the same chromosome that are lost in SMCHD1 knockout cells (Fig. 4A, Fig. 4C-D; Extended Data Fig. 6A and 6B). This data supports the proposal that SMCHD1 is an important anchor point for heterochromatin regions that are distant from each other on the linear chromosomes. A more high-resolution analysis shows that there are many contact points (dots) between two individual B compartments (B-to-B contacts) as indicated by a cloud of dots in shapes of distinct broader stripes (Fig. 4A; Extended Data Fig. 6A, 6B). However, these dots and stripes disappear in the SMCHD1 knockout cells.

**Figure 4:**
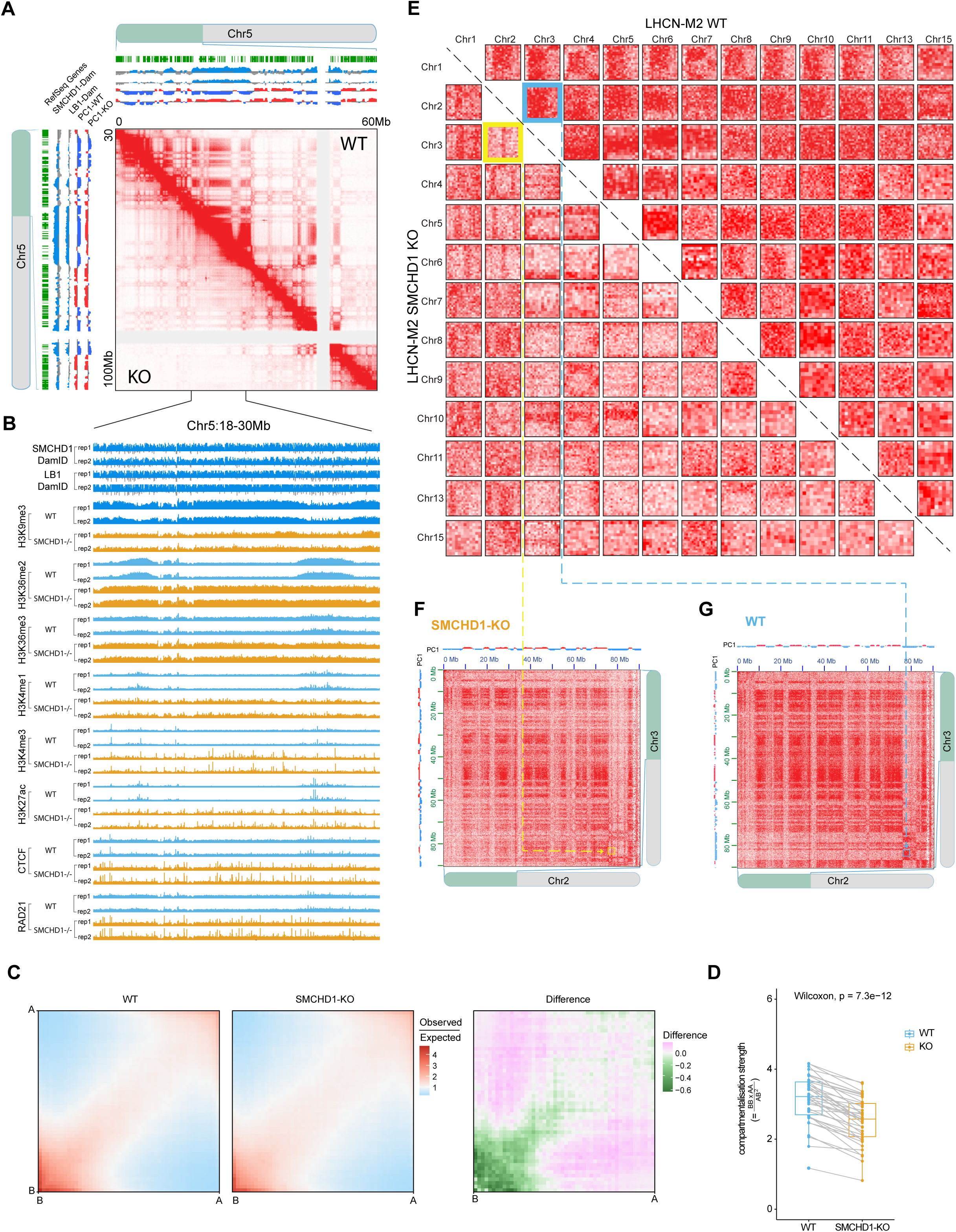
Loss of contacts between B compartments in SMCHD1-depleted cells. **A**. In situ Hi-C contact map showing contacts between individual B compartments (broad stripes) across a 60-Mb region of chromosome 5 and their loss in SMCHD1^−/−^ cells. These contacts are not found in the absence of SMCHD1. The SMCHD1 and Lamin B1 (LB1) signals are shown along with B compartments (blue) and A compartments (red) on top and left. **B.** SMCHD1, Lamin B1 (LB1) and histone modification patterns in WT and SMCHD1^−/−^ cells in a magnified representative region from panel A. **C.** Saddle plots showing the genome-wide compartmentalization between WT and SMCHD1^−/−^ cells. Differential saddles with green colors denoting fewer B-to-B inter-compartment interactions in SMCHD1^−/−^ cells compared to wildtype cells are shown on right. **D.** Boxplot of the compartmentalization strength per each chromosome arm (dots). This data corresponds to the proportion of inter-compartment (B-to-B and A-to-A) contacts versus inter-compartment (A-to-B and B-to-A) contacts and is calculated for each chromosome arm separately. Paired Wilcoxon test was used for statistical analysis. **E.** Loss of interchromosomal B-to-B compartment contacts seen between 156 pairs of the longer chromosomes. Each square shows a representative interchromosomal B-to-B compartment contact region in WT cells (top right) that is lost in SMCHD1-KO cells (bottom left). **F.** Expanded view of contact maps between chr2 and chr3 (showing the position of the yellow square from panel E for the SMCHD1-KO. A-compartments (red) and B-compartments (blue) are shown along the chromosomes. **G.** Expanded view of the blue square B-to-B compartment region shown in panel E for WT cells.

The B-to-A compartment transitions and loss of inter-B compartment contacts are associated with loss of the repressive histone mark H3K9me3 and gain of many active marks, including H3K36me2, H3K36me3, H3K4me1, H3K4me3, H3K27Ac, as well as gain of CTCF and RAD21 binding peaks (Fig. 4B; Extended Data Fig. 6C, 6D, 6E). Figure 4C shows the loss of B-to-B compartment contacts in SMCHD1^−/−^ cells using genome-wide analysis. Compartmentalization strength (inter-B and inter-A) is decreased along all chromosome arms except for chr 19q, a very gene-rich chromosome arm, which shows a minor reduction (Fig. 4D).

In addition to observing the loss of B-to-B compartment contacts on the same chromosome in absence of SMCHD1, our high sequencing depth allowed us to examine interchromosomal contacts in many pairwise combinations (Fig. 4E-G). The strongest contact densities between different chromosomes were observed for A-to-A compartment contacts (see for example Fig. 4F, 4G). However, almost every chromosome contained one or more strong signal for B-to-B contact regions with another chromosome in wildtype cells that were strongly diminished in the SMCHD1-deficient cells. Examples are shown in Figure 4E for 156 of such lost interchromosomal B-B contact regions. Zoom-out views are shown for a part of chromosomes 2 and 3 (Fig. 4F, 4G). The data shows that SMCHD1 is not only critical for joining B compartments on the same chromosome but also maintains such compartment connections between different chromosomes.

### SMCHD1 organizes LADs and 3D genome structure

B to A compartment transitions were targeted to SMCHD1-bound regions (Fig. 3). We observed that these same regions lose their association with Lamin-B1 as shown by diminishing Lamin B1 signal over B-to-A transitioning compartments (Fig. 5A, 5B). To further analyze this phenomenon at the global level, we performed Chrom3D, a method that uses 3D TAD information obtained from the HiC data along with data on Lamin-B1-associated regions to construct a 3D model of nuclear and chromatin structure ^43,44^. Analyzing the distribution of the chromosomes inside the nucleus, we observed a striking shrinkage of the territorial volume that the chromosomes occupy in the SMCHD1-deficient cells relative to wildtype cells (Fig. 5C; Movies S1 and S2). Chrom3D shows the peripheral localization of LADs in the nuclear shell in wildtype myoblasts (Fig. 5D; see Movies S3 and S5). However, in SMCHD1-KO cells, much of the Lamin B1 signal has disappeared from the nuclear periphery and shows a more random organization (Fig. 5D; Movies S4 and S6).

**Figure 5:**
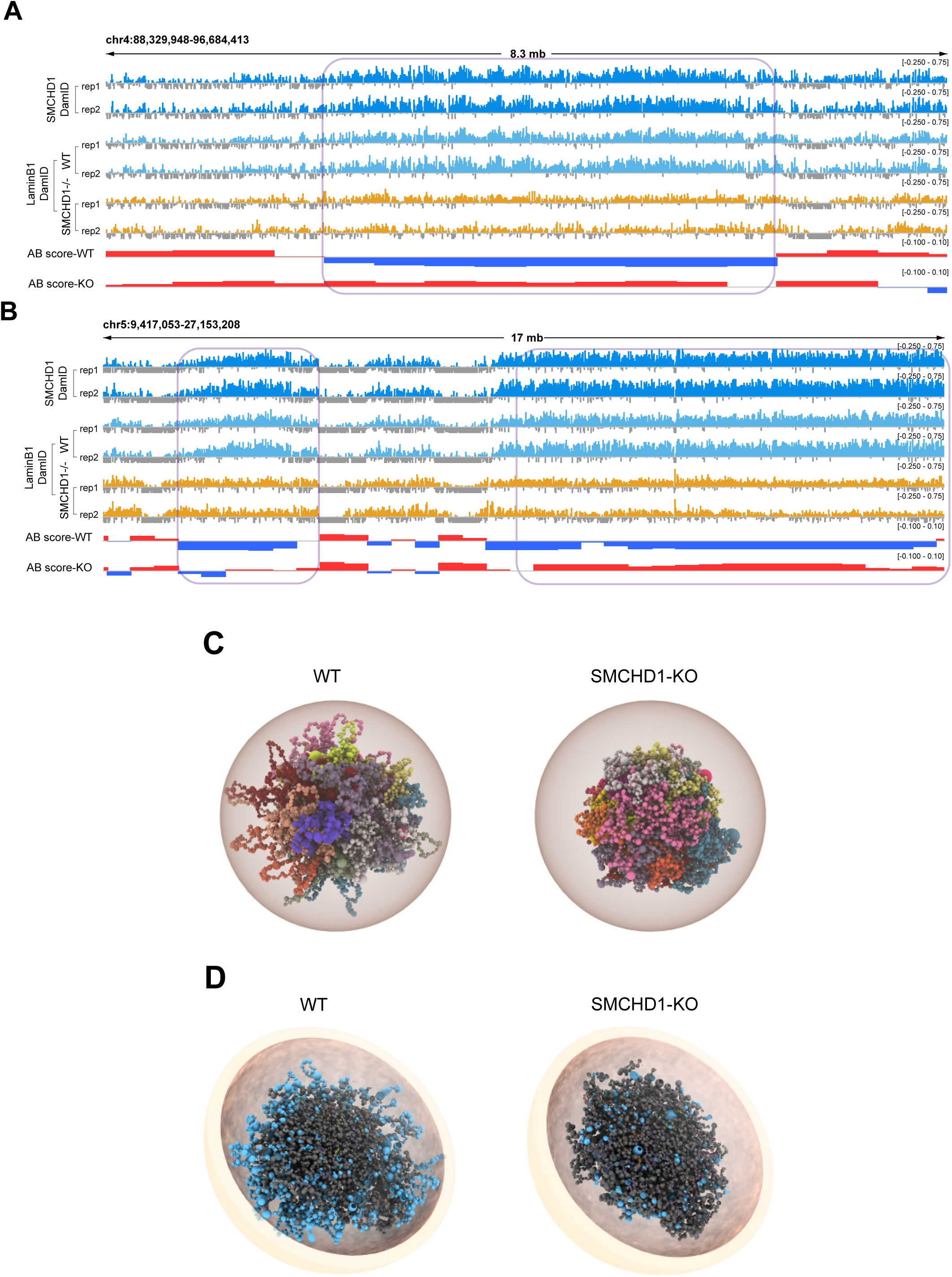
Loss of LADs and alterations of 3D chromatin structure in SMCHD1-deficent cells. **A**. Plots showing changes in Lamin B1 over a B-to-A compartment transitions for a representative region on chr4 in WT and SMCHD1^−/−^ cells. Two replicates each of Dam-ID sequencing are shown along with B compartments (blue) and A compartments (red). The signals were normalized relative to Dam-only signal. **B.** Genome browser views of an area of chr5 showing the distribution of SMCH1 and of Lamin B1 in wildtype and in SMCHD1-KO cells (see panel A for details). **C.** 3D structure showing spatial distribution of individual chromosomes for the whole genome in WT and SMCHD1-KO cells. Radius is 5 μm. **D.** Cross-sectioned view of 3D structure showing spatial distribution of individual chromosomes for the whole genome in WT and SMCHD1-KO cells. The LAD-associated genomic regions are indicated as blue spheres. Grey spheres represent TADs without LAD association. Radius is 5 μm.

At the level of single chromosomes, we noted an outside localization of the Lamin B1 associated regions on chromosomes 2, 3, and 6 (and most other chromosomes, not shown) in wildtype cells but a more compact organization of the chromosomes in the SMCHD1-deficient cells (Extended Data Fig. 7A-C). The exceptions are smaller, gene-rich chromosomes, which have few LADs and few SMCHD1-bound sequences. An example is chromosome 19, where substantial Lamin B1 signals are only found near the centromere (Extended Data Fig. 7D). This chromosome shows no major structural differences between wildtype and knockout cells (Extended Data Fig. 7D) suggesting that the structural changes we observe on the other chromosomes are linked to SMCHD1/Lamin B1-bound genomic features. Combined, the data shows that SMCHD1 is an important regulator that maintains genome compartments and nuclear structure.

### Topologically associated domains and loops are remodeled by inactivation of SMCHD1

When zooming in to reveal further details, in parallel with the B-to-A compartment transitions and the loss of the long-range contacts between different B compartments, we observe a gain of distinct sets of contacts suggesting the emergence of new topologically associated domains (TADs) (Fig. 6A, 6B; Extended Data Fig. 8A, 8B). Aligning this data with ChIP-seq peaks of CTCF and RAD21 shows that the new TAD boundaries are delineated by new CTCF and RAD21 binding (Fig. 6C, 6D; Extended Data Fig. 8C). Heatmap and histogram profile of insulation scores showed strong insulation at the CTCF and RAD21 binding sites in SMCHD1^−/−^ cells at a global level, compared WT cells (Fig. 6C, 6D). Aggregate TAD analysis is depicted in Extended Data Fig. 8D and shows an increase in TAD formation upon loss of SMCHD1, both within TADs and with neighboring TADs (Extended Data Fig. 8E). Furthermore, cumulative distribution plots (CDPs) for insulation scores comparing WT versus SMCHD1^−/−^ cells also showed a statistically significant difference for the whole genome (P<2.2e-16), with the slope of the SMCHD1^−/−^ curve being shallower before zero and steeper after zero, indicative of greater fluctuation of insulation scores and more TADs (Extended Data Fig. 8F). A similar pattern was also observed on presentative chromosomes 4, 10 and 13 in WT and SMCHD1^−/−^ cells (Extended Data Fig. 8G).

**Figure 6:**
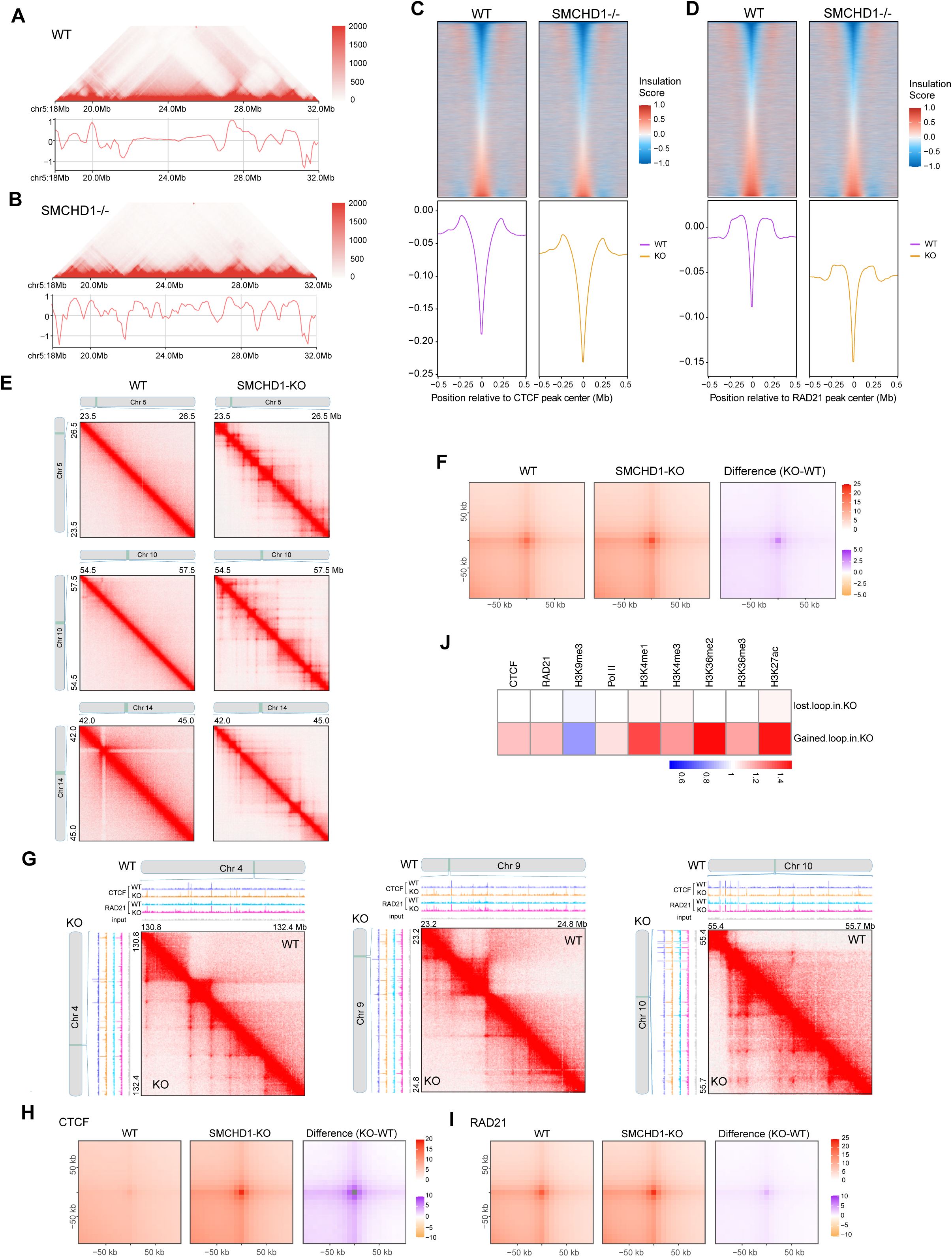
Gains of TADs and loops in SMCHD1-deficient cells are linked to CTCF and Cohesin/RAD21 binding. **A**. Hi-C contact map and insulation profiles at a representative region on chromosome 5 in wildtype cells. **B.** Hi-C contact map and insulation profiles at a representative region on chromosome 5 in SMCHD1^−/−^ cells. **C.** Insulation heatmap and histogram profile of insulation scores at 10 kb resolution spanning a 500 kb window at CTCF peak centers between WT and SMCHD1^−/−^ cells. The color maps show strong insulation in blue and weak insulation in red. **D.** Insulation heatmap and histogram profile of insulation scores at 10 kb resolutions spanning a 500 kb window at RAD21 peak centers between WT and SMCHD1^−/−^ cells. Color maps show strong in blue and weak insulation in red. **E.** Hi-C contact maps showing more loops at representative regions on chromosomes 5, 10, and 14 after loss of SMCHD1. **F.** Aggregate peak analysis (APA) of all chromatin loops detected in wild-type and SMCHD1^−/−^ cells, which is showing increased interaction intensity in SMCHD1^−/−^ cells versus wild-type cells. The differential APA analysis plot with purple denotes more interactions in SMCHD1^−/−^ cells compared to the wildtype cells. **G.** Gains of loops at representative regions on chromosomes 4, 9, and 10, linked to newly arising RAD21 and CTCF peaks in SMCHD1 KO cells as shown in the ChIP-seq tracks. **H.** APA analysis of chromatin loops located at CTCF binding sites detected in wildtype and SMCHD1^−/−^ cells, which shows increased interaction intensity in SMCHD1^−/−^ cells versus wild-type cells. Differential APA analysis plot with purple denotes more interactions in SMCHD1^−/−^ cells compared to wildtype cells. **I.** APA analysis of chromatin loops located RAD21 binding sites detected in wildtype and SMCHD1^−/−^ cells, which shows increased interaction intensity in SMCHD1^−/−^ cells versus wild-type cells. A differential APA analysis plot with purple denotes more interactions in SMCHD1^−/−^ cells compared to the wildtype cells. **J.** Heatmap showing differences in CTCF, RAD21, Pol II, and chromatin marks over gained or lost loop intervals in SMCHD1^−/−^ cells.

Our dataset provided sufficient resolution to examine loop formation well below the 5kb resolution. We find that the loss of SMCHD1 leads to the formation of many new loops (e.g., dots in Fig. 6E, 6G; Extended Data Fig. 8C). At the whole genome level, we saw an increase of contacts at the loops in the absence of SMCHD1 (Fig. 6F). The new loops are anchored at newly arising CTCF and RAD21 peaks in the SMCHD1 knockout cells, in some cases in combination with CTCF peaks that preexisted in the wildtype cells (Fig. 6G). Global analysis of chromatin loops located at CTCF or RAD21 binding sites detected in wildtype and SMCHD1^−/−^ cells shows increased interaction intensity in SMCHD1^−/−^ cells (Fig. 6H, 6I). The new loop formation was associated with the gain of active chromatin components, most strongly with gains of H3K4me1, H3K36me2, and H3K27Ac and with a loss of H3K9me3 (Fig. 6J).

### SMCHD1 regulation of enhancer-promoter interactions and transcription

Changes in gene expression in the knockout cells were relatively limited, likely reflecting the fact that SMCHD1 chiefly is associated with gene-poor regions of chromosomes. Using RNA-seq, we found 208 upregulated genes and 180 downregulated genes (fold change >2, FDR<0.05) (Fig. 7A; Extended Data Table 3). Gene ontology analysis shows enrichment of terms related to muscle cell development and neurogenesis (Fig. 7B; Extended Data Fig. 9A).

**Figure 7:**
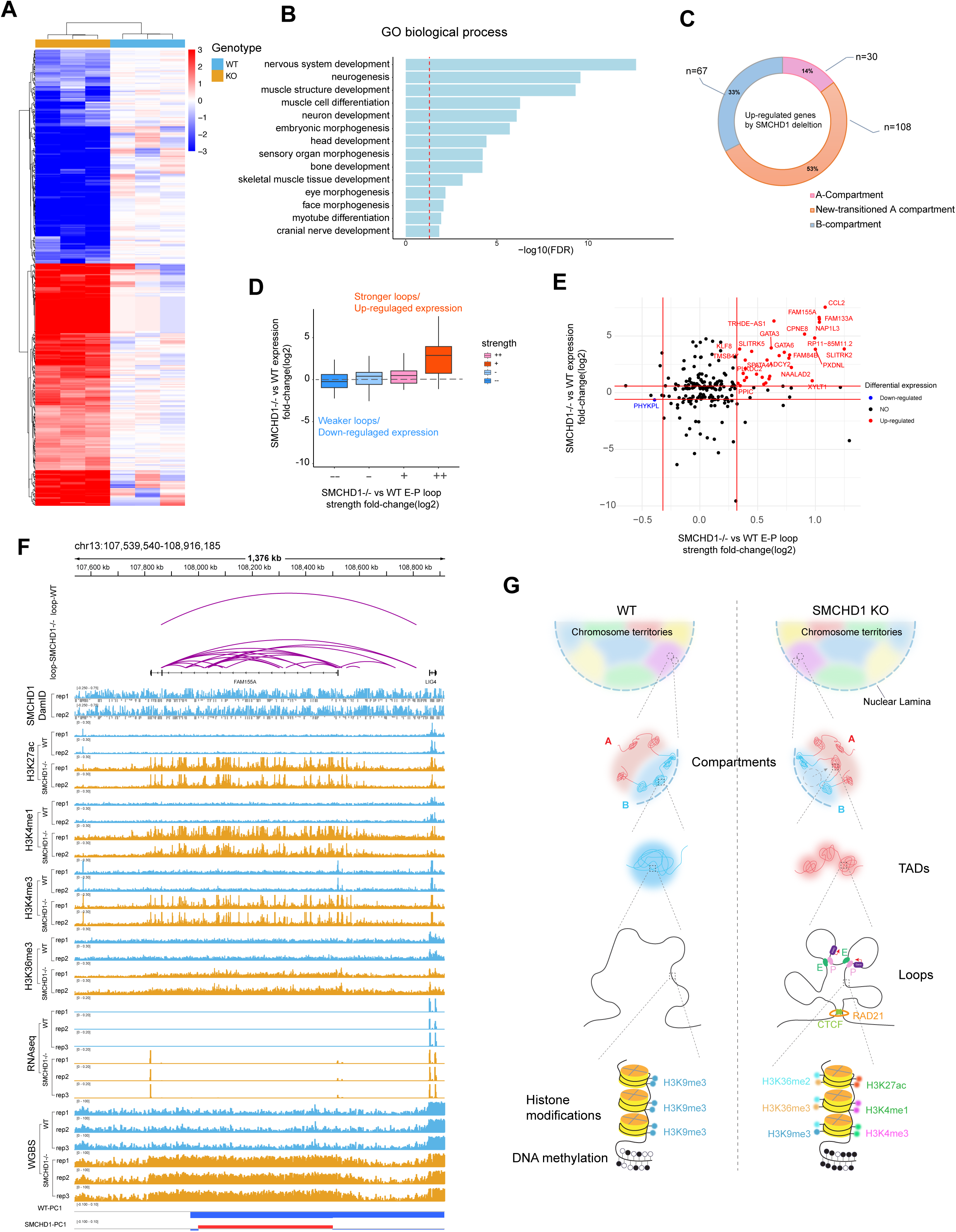
Changes in gene expression and enhancer-promoter interactions after loss of SMCHD1. **A**. Heat map of gene expression changes. **B.** GO analysis of differentially expressed genes. **C.** The relationship between differential gene activation and compartment transitions. **D.** Boxplots depicting expression fold change (log2) between SMCHD1^−/−^ and WT cells (y axis) for genes engaged in enhancer-promoter (E-P) loops. Genes are stratified by change in E-P loop strength between WT and SMCHD1^−/−^ cells (x axis). **E.** Scatter plot presenting the relationship between gene expression and enhancer-promoter (EP) loop strength between WT and SMCHD1^−/−^ cells. **F.** The *FAM155A* gene is upregulated in parallel with acquisition of active marks and extensive new loop formation in SMCHD1^−/−^ cells. This panel shows browser tracks for RNA-seq data, distribution of active histone marks and mapped loops in WT and SMCHD1 KO myoblasts. **G.** Model of SMCHD1 function in 3D epigenome landscape maintenance.

Of particular interest were those upregulated genes that we found in SMCHD1-bound regions which underwent B-to-A compartment transitions, usually in parallel with acquisition of active chromatin marks. More than half of all upregulated genes were localized within B-to-A transitioning compartments (Fig. 7C). We next determined if formation of new enhancer-promoter (EP) contacts may globally underlie gene activation events in SMCHD1-deficient cells. Figures 7D and 7E show that higher gene activation levels paralleled the extent of new EP contact formation. Examples are shown in Extended Data Fig. 9B for the genes *CCL2* and *NAP1L3*. These de novo loop formations are associated with gain of active chromatin marks (Extended Data Fig. 9B, 9C). Upregulated genes are associated with gain of active marks whereas downregulated genes loose active marks, as expected (Extended Data Fig. 9D).

The likely disease-causing gene activated in muscle cells of FSHD2 patients that carry SMCHD1 mutations and/or a contraction of the D4Z4 repeat units at chromosome 4q35 (FSHD1) is *DUX4* ^18^ encoding an embryonic transcription factor ^45–47^. Using our RNA-seq data and RT-PCR, we did not observe an upregulation of *DUX4* expression (Extended Data Fig. 9E, Extended Data Fig. 10A), nor did we observe upregulation of DUX4 target genes in SMCHD1 knockout cells (Extended Data Fig. 9E). In addition, DNA methylation patterns at the telomeric *DUX4* repeat unit did not change after SMCHD1 loss (Extended Data Fig. 10B). This finding is consistent with previous results in which inactivation of SMCHD1 in human somatic cells did not reactivate this locus ^38^. We were only able to activate *DUX4* and demethylate the locus by combined inactivation of SMCHD1 and treatment of the cells with the DNA methylation inhibitor 5-azadeoxycytidine (5-azaC-dR) but not with 5-azadeoxycytidine alone (Extended Data Fig. 10C, 10D).

Some of the most highly upregulated genes are indicated in Figure 7E. One of these genes is *FAM155A*, encoding a component of hetero-tetrameric NALCN sodium channels ^48–50^. *FAM155A* was highly upregulated after SMCHD1 knockout (Fig. 7E, 7F). At the *FAM155A* locus, we observed de novo formation of CTCF and RAD21 binding sites and extensive de novo loop formation in absence of SMCHD1 along with gain of many active chromatin modifications (Fig. 7F). In summary, the loss of SMCHD1-regulated chromatin organization leads to perturbation of gene expression programs in the affected cells with a potential contribution to the etiology of diseases with dysfunctional SMCHD1 protein.

## DISCUSSION

SMCHD1 is mutated in the human muscular dystrophy, FSHD2, which is characterized by activation of the *DUX4* homeobox gene ^18^. We did not observe activation of *DUX4* gene in somatic muscle cells lacking SMCHD1, except when the cells were additionally exposed to a DNA methylation inhibitor. Dysfunction of SMCHD1 during developmental time windows when gene repression is established at the *DUX4* locus may be a critical event in disease initiation. On the other hand, the gene expression changes we did observe in myoblasts lacking functional SMCHD1 may contribute to FSHD2. One of the most strongly upregulated genes was *FAM155A*, a component of sodium channels. Alterations in the NALCN sodium leak channels, which FAM155A is a component of, have been linked to hypotonia, a form of muscle weakness ^51^. It is conceivable that an imbalance of NALCN subunits due to strong overexpression of *FAM155A* may contribute to a muscle cell phenotype. It remains to be determined if and how changes to the nuclear structure as we observed in SMCHD1-deficient cells may also contribute to FSHD. Myocytes utilize nuclear mechanosignaling mechanisms to respond to changes in the environment and transmit them through the nuclear envelope and lamina to the nucleus to alter transcription. Intriguingly, mutations in several nuclear envelope and lamina-associated proteins, Lamin A/C, Emerin, Nesprin1/2, and LAP1, are all linked to different forms of muscular dystrophy ^52,53^.

Here we present the first genome-wide analysis of SMCHD1 function in human muscle cells. We show that SMCHD1 colocalizes and copurifies with Lamin B1 and is colocalized extensively with the LAD-associated heterochromatin mark H3K9me3. Depletion of SMCHD1 leads to a reduction of H3K9me3 along many SMCHD1-bound regions, a visible loss of heterochromatin at the nuclear periphery as determined by transmission electron microscopy, and to a loss of LADs. Unexpectedly, the loss of H3K9me3 occurs in parallel with a gain of DNA methylation which increases over megabase size regions that loose H3K9me3. The loss of SMCHD1 leads to an uncoupling of the two marks.

Along with the loss of H3K9me3, we observe a gain of many active chromatin marks, including promoter and enhancer marks (H3K4me3, H3K4me1, H3K27Ac), active gene body marks (H3K36me3, RNA polymerase), the more broadly distributed euchromatin modification H3K36me2, as well as the chromosomal structural proteins, CTCF and RAD21. The changes observed reflect the conversion of heterochromatin to euchromatin (B-to-A compartment transitions) at SMCHD1-bound regions as determined by 3D compartment analysis. In this context, it is of note that DNA CpG methylation, although commonly associated with gene repression when it occurs at promoters or enhancers, should also be viewed as a euchromatin mark. DNA methylation levels are much higher in gene-dense regions relative to gene-poor regions of chromosomes. DNA methylation in gene bodies has an activating role in gene regulation and is positively correlated with gene expression states ^54,55^.

The fact that SMCHD1 plays an important part in organizing large stretches of constitutive heterochromatin on autosomes in human somatic cells has previously not been appreciated. Prior studies have focused predominantly on the function of SMCHD1 in X chromosome inactivation ^14,21–24^ or in the control of a few select autosomal gene clusters, such as the *HOX* genes ^31^. Although the structure of the inactive X chromosome is unique, there are some similarities of SMCHD1 function in X inactivation and in autosomal heterochromatin maintenance. SMCHD1 antagonizes TADs and compartmentalization and merges the S1 and S2 compartments on the inactive X chromosome in mouse embryo fibroblasts or mouse neural progenitor cells (Xi) ^21–23^. However, a role of H3K9me3 in X chromosome inactivation is less defined. SMCHD1 was shown to be recruited to the Xi by a pathway involving XIST, HNRNPK and the PRC1 Polycomb complex via histone H2A K119 ubiquitylation ^25^. Polycomb components play an important role in X chromosome inactivation ^56,57^ but their contribution to constitutive heterochromatin is not clear since this mark occurs across both A and B compartments. Furthermore, SMCHD1 is critical for XIST RNA spreading on the Xi ^24^. The exact roles of SMCHD1 on the inactive X and at autosomal heterochromatin regions are expected to be different. Prior Hi-C studies of SMCHD1 knockouts have not observed substantial compartment switching or changes in TADs on autosomes ^21–23^. However, Wang et al. did find that SMCHD1 was localized in gene-poor regions of mouse embryo fibroblast cells ^23^, where the protein colocalizes with lamina-associated domains ^58^. In a proteomics study, SMCHD1 was found as an interactor of the inner nuclear membrane bound lamina-associated protein LAP2β, along with the proteins HP1 β, LRIF1, and EHMT2, a H3K9 methyltransferase ^58^. These unexpected differences of our data from published work raise the important questions as to whether our finding is unique to muscle cells, occurs only in human and not in mouse cells (unlikely, given the inter-species conservation of the SMCHD1 protein), or is a feature of only male cells, which lack an inactive X chromosome. Future work should address all these possibilities.

Removal of SMCHD1 leads to the loss of many inter-B compartment contacts, along the same or between different chromosomes, a weakening of B compartments and frequent switches from B into A compartments. Thus, the SMCHD1 protein functions as an important global tether of heterochromatic B compartments, likely by tying a substantial fraction of heterochromatin to the nuclear lamina (see model in Fig. 7G).

Our data suggests that the B compartment (heterochromatin) is physically inaccessible in wildtype cells to a group of euchromatic histone and DNA modification enzymes and to at least some structural proteins. The model is consistent with a study in which promoters from numerous silent LAD-associated genes show strong activity when moved onto episomal plasmids^34^. One particularly surprising finding concerns DNA methylation changes which occurred as long stretches of hypermethylation within the B-to-A switched regions in absence of SMCHD1. The hypermethylation is likely carried out by the de novo DNA methyltransferases DNMT3A and/or DNMT3B. The activities of these enzymes are stimulated by the euchromatic histone marks H3K36me2 for DNMT3A ^59^, and by H3K36me3 for DNMT3A and DNMT3B ^54,60^. These two histone modifications also substantially increase within the B-to-A converted regions (Fig. 4B, Extended Data Fig. 5C, 6E). It is a general misconception that heterochromatin is enriched in DNA methylation. The partial methylation of LADs has typically been explained by late replication or a DNMT sequence preference for these domains ^61,62^. However, here we offer another explanation, namely that there is much-reduced accessibility of heterochromatin to the de novo DNA methyltransferases and the H3K36 histone methyltransferases which create improved substrates for the DNA methylation enzymes. These regions become easily H3K36-methylated, and de novo DNA-methylated upon a compartment switch (B-to-A). We propose that A compartments are formed by default when not prevented by SMCHD1.

We report here that regions of gene-poor heterochromatin will flip to become euchromatin by removal of a single chromosomal structural protein. Our data suggests that the spatial segregation of chromatin drives its transcriptional state rather than resulting from it. The 3D genome architecture at the sub-megabase scale is regulated by the CTCF/Cohesin system, in which loop extrusion by Cohesin and the arrest of this process at CTCF bound sites leads to the formation of loops and TADs. Notably, depletion of CTCF causes the disappearance of most TADs while leaving the higher order organization of chromatin into A and B compartments largely unchanged ^63^. The cellular determinants of compartment organization have remained elusive. Here we report that the protein SMCHD1 is a critical component of the molecular machinery that maintains genome compartments, 3D-chromatin architecture, and epigenome landscape.

## ACKNOWLEDGEMENTS

We thank the members of the Van Andel Institute Core Facilities (Genomics, Bioinformatics and Biostatistics, Optical Imaging, Cryo-electron Microscopy) for their support and technical assistance.

## Funding

This work was supported by NIH grant R01AR079174 to GPP.

## Author contributions

Conceptualization, G.P.P., Z.H.; experimental design, Z.H., W.C., and G.P.P.; investigation, Z.H., W.C., I.R., R.T.; data analysis, Z.H., W.C., and G.P.P.; writing, Z.H. and G.P.P.; visualization, Z.H., I.R.; supervision, G.P.P.; funding acquisition, G.P.P.

## Competing of interests

The authors declare no conflicts of interest.

## Data and materials availability

All data are available in the main text or the Supplementary Material.

Raw sequencing files and key intermediate files generated in this study are deposited and freely available from Gene Expression Omnibus (GEO: GSE251747-GSE251751). All data is provided at GEO GSE251747 (ChIP-seq, reviewer token yhapecwoxpwlvar), GSE251748 (Dam-ID-seq, reviewer token ydelsywcdtgbfid), GSE251749 (HiC, reviewer token mbszwuekznoxten), GSE251750 (RNA-seq, reviewer token mdifsuigrxwdtyt), and GSE251751 (WGBS, reviewer token erwxwmcinzarzqv).

## List of Supplementary Materials

Materials and Methods

Extended Data Figures 1 to 10

Extended Data Tables 1 to 3

Movie S1 to S6

## SUPPLEMENTARY MATERIAL

### Materials and Methods

#### Cell lines

LHCN-M2, a human myoblast cell line immortalized with hTERT and CDK4, was obtained from Evercyte (Vienna, Austria). LHCN-M2 cells were cultured on 0.1% gelatin-coated tissue culture plates in DMEM / medium 199 (4:1) supplemented with 15% fetal bovine serum, 0.02 M HEPES, pH 7.1, 0.03 μg/mL zinc sulfate, 1.4 μg/mL vitamin B12, 0.055 μg/mL dexamethasone, 2.5 ng/mL hepatocyte growth factor, 10 ng/mL basic FGF, 60 units/mL penicillin and 60 μg/mL streptomycin.

Myoblasts from healthy individuals (n = 3) or from patients with FSHD2 (n = 3) were obtained from the FSHD Registry at the University of Rochester. The genotype of 19MB015 was SMCHD1+: c.610A>G, that of 19MB016 was SMCHD1+: c.1647+3A>G, and 19MB017 had a 1.2 megabase deletion encompassing the *SMCHD1* gene. Normal and patient myoblasts were cultured in F-10 Nutrient Media supplemented with 20% FBS, 1% penicillin/streptomycin, 10 ng/ml bFGF and 1 µM dexamethasone.

#### Generation of knockout LHCN-M2 myoblast lines using CRISPR-Cas9

SMCHD1^−/−^ LHCN-M2 cell lines were generated by pLentiCRISPR-E with the blasticidin resistance gene generated from pLentiCRISPR-E-puromycin (a gift from Phillip Abbosh; Addgene plasmid # 78852). The pLentiCRISPR-E-Blast vector carrying the appropriate SMCHD1 sgRNAs was co-transfected into HEK293FT cells with lentiviral packing plasmid psPAX2 and envelope plasmid pMD2.GVG. For transfection of a 10 cm dish, FuGENE HD reagent (Promega, E2311) was diluted into 2 ml Opti-MEM and then the following DNA was added: 9.9 μg plentiCRISPR-E-Blast, 7.5 μg psPAX2 and 5 μg pMD2.GVG. Viral particles were harvested at 48 h after transfection and frozen at –80°C. LHCN-M2 cells were transduced with plentiCRISPR-E-Blast-SMCHD1 virus, and single-cell clones were selected in 10 μg/mL blasticidin (10 μg/mL). Protein expression levels in the knockout cells were confirmed by Western blot. All SMCHD1 knockout clones were further confirmed by the presence of frameshift mutations detected by Sanger sequencing of the CRISPR-targeted region, which was PCR-amplified from genomic DNA and cloned into Topo-TA cloning vector (Thermo, 450030).

#### Western blot

Western blots were performed as previously described ^30^, with minor modifications. Cells were lysed in lysis buffer (50 mM Tris-HCl, pH 7.4, 150 mM NaCl, 1% Triton X-100, 1 mM EDTA and proteinase inhibitor cocktail (Roche,11873580001) on ice for 1 hour, then the cell lysate was centrifuged at 12,000xg for 15 min at 4°C. The centrifuged cell lysates were then separated on 4%-15% SDS-polyacrylamide gels and transferred onto PVDF membranes (Bio-Rad) by wet transfer at 4°C. The membranes were incubated with blocking buffer (5% non-fat milk, 0.1% Tween-20 in PBS) for 1 hour at room temperature, and the membranes were then incubated with the indicated primary antibody at 4°C overnight. We washed the PVDF membranes with PBS-Tween (0.1%), followed by incubation with peroxidase-conjugated secondary antibodies for 1 hour at room temperature. Blotting signals were detected using ECL Prime detection reagent (GE Healthcare). Antibodies used for Western blots were anti-SMCHD1 (1:2,500, Bethyl Laboratories, A302-871A), anti-alpha-tubulin (1:10,000, Abcam, ab7291), HRP goat anti-rabbit IgG (Active Motif, 1:10,000, 15015), HRP goat anti-mouse IgG (1:10,000, Active Motif, 15014). Lamin B1 (Abcam, ab16048), G9a/EHMT2 (R&D Systems, PP-A8620A-00), SETDB1 (Proteintech, 11231-1-AP), H3K9me3 (Abcam, ab8898) and histone H3 (Abcam, ab1791).

#### Chromatin fractionation

Chromatin fractionation was performed as previously described ^63^, with minor modifications. LHCN-M2 cells were washed in PBS and extracted in cytoskeleton (CSK) buffer (10 mM PIPES, pH 6.8, 100 mM NaCl, 1 mM EGTA, 300 mM sucrose, 3 mM MgCl_2_, protease inhibitors, and 1 mM phenylmethylsulfonyl fluoride (Thermo Fisher scientific, 36978) supplemented with 1 mM dithiothreitol and 0.5% Triton X-100). We incubated the reaction on ice for 5 mins, and then separated the cytoskeletal fraction from soluble proteins by centrifugation at 800xg for 3 min. The supernatant was designated as the S1 fraction. The pellets were washed with an additional volume of CSK buffer. We solubilized chromatin by adding 25 U of ribonuclease-free DNase (Invitrogen) in CSK buffer at 37°C for 30 min, followed by adding ammonium sulfate in CSK buffer to a final concentration of 250 mM, then incubated the reaction on ice for 5 mins, and pelleted the samples at 2,500xg for 3 min at 4°C. The supernatant was designated as the S2 fraction. We washed the pellet with CSK buffer, then treated the pellet with CSK buffer including 2 M NaCl for 5 min at 4°C and centrifuged at 2,500xg for 3 min at 4°C. The supernatant after this step was designated as the S3 fraction. We washed the remaining insoluble pellet with CSK buffer supplemented with 2 M NaCl twice. The insoluble pellets were then treated with 8 M urea buffer and the solubilized sample was considered as the nuclear matrix–containing fraction (S4). Supernatants from each extraction step were quantified and analyzed by SDS-PAGE and immunoblotting.

#### RNA preparation and RNA sequencing (RNA-seq)

Total RNA was extracted from LHCN-M2 cells using the PureLinkTM RNA Mini kit (Ambion, 12183020), according to the manufacturer’s protocol. Total RNA integrity was verified by Agilent 2100 Bioanalyzer (Agilent Technologies) and quantified with a NanoDrop 8000 instrument (Thermo Fisher). RNA-seq libraries were prepared from total RNA with the Standard Kapa stranded mRNA library prep Kit (KAPA Biosystems, KR0960) according to the manufacturer’s protocols. RNA-seq was performed with three biological replicates per LHCN-M2 muscle cell lines (WT and SMCHD1 KO). Library size distributions were validated on the Bioanalyzer (Agilent Technologies). Sequencing was performed with an Illumina NextSeq500. Library de-multiplexing was performed following Illumina standards.

#### Whole-genome bisulfite library preparation and sequencing

The total genomic DNA was isolated using the Quick-DNA Miniprep Plus kit (Zymo Research, D4070). WGBS libraries were prepared according to the manufacturer’s instructions using the Swift, Accel-NGS Methyl-Seq DNA Library Kit (Swift Biosciences, 30024) and Zymo’s EZ DNA Methylation-Lightning kit (Zymo Research, D5030). Sequencing was performed with an Illumina Novaseq 6000 instrument with 150-bp paired-end read runs.

#### ChIP-seq

Chromatin state maps (H3K27ac, H3K4me1, H3K4me3, H3K36me2, H3K36me3 and H3K9me3) and binding profiles for RNA polymerase II, CTCF insulator protein and the RAD21 cohesin protein subunit were generated by ChIP-seq. Briefly, cells were cross-linked with 1% formaldehyde for 10 min at room temperature and the reaction was quenched with 125 mM glycine. Crosslinked cells were lysed in lysis buffer (50 mM HEPES-KOH, pH 7.9, 140 mM NaCl, 1 mM EDTA, 10% glycerol, 0.5% NP40, 0.25% Triton X-100, supplemented with cOmplete protease inhibitors (Roche)) and incubated on ice for 10 min. Following centrifugation at 4,500 rpm for 5 mins at 4°C in a benchtop centrifuge, the cell pellets were then washed with wash buffer (10 mM Tris-Cl, pH 8.1, 200 mM NaCl, 1 mM EDTA, pH 8.0, 0.5 mM EGTA, pH 8.0, supplemented with cOmplete proteinase inhibitors) and shearing buffer (0.1% SDS, 1 mM EDTA, 10 mM Tris-Cl, pH 8.1, supplemented with cOmplete proteinase inhibitors). Then, the cell pellets were resuspended in shearing buffer and sonicated with a Covaris E220 Evo sonicator to shear DNA to 300 to 500 bp size fragments. The resulting lysate was supplied with NaCl and Triton X-100 to reach a final concentration of 150 mM NaCl and 1% Triton X-100, then cleared by centrifugation for 10 min at 20,000xg, and then incubated with washed Dynabeads Protein G (Invitrogen, 10004D) and antibody overnight on a rotator at 4°C. Antibodies were as follows: CTCF (Active Motif, 61311), H3K27ac (Abcam, ab4729), H3K9me3 (Abcam, ab8898), H3K36me2 (CST, 2901S), H3K36me3 (Abcam, ab9050), H3K4me1 (Abcam, ab8895), H3K4me3 (Millipore, 07-473), Pol2 (Millipore, 05-623) and RAD21 (Abcam, ab992).

Beads were then collected and washed with low salt buffer (0.1% SDS, 1% Triton X-100, 2 mM EDTA, 20 mM HEPES-KOH, pH 7.9, 150 mM NaCl supplemented with cOmplete proteinase inhibitors), high salt buffer (0.1% SDS, 1% Triton X-100, 2 mM EDTA, 20 mM HEPES-KOH, pH 7.9, 500 mM NaCl supplemented with cOmplete proteinase inhibitors) and LiCl buffer (100 mM Tris-Cl, pH 7.5, 500 mM LiCl, 1% NP40, 1% sodium deoxycholate) twice and with TE buffer (10 M Tris-Cl, pH 8.1, 1 mM EDTA) once. ChIP DNA samples were eluted with Proteinase K digestion buffer (20 mM HEPES, pH 7.9, 1 mM EDTA, 0.5% SDS) and 1 µl of Proteinase K (20 mg/mL). Purified DNA was quantified for library preparation with Qubit sensitivity dsDNA HS Kit. Libraries were then prepared using the TruSeq ChIP Sample Preparation Kit (Illumina, IP-202–1012, IP-202–1024) according to the manufacturer’s instruction. Briefly, 5 ng of ChIP-DNA was used for input and IP samples. Libraries were amplified using 14 cycles on a thermocycler. Libraries then were quantified and validated using the Agilent High Sensitivity DNA Kit and bioanalyzer. Sequencing was performed with an Illumina HiSeq 2500 instrument with 150-bp paired-end read runs. All ChIP-Seq experiments were processed in parallel with whole cell extract input controls.

#### DamID-sequencing and library preparation for SMCHD1 and Lamin B1

To map the enrichment patterns of SMCHD1 and Lamin B1, we generated DamID-seq libraries following a protocol described previously ^23,64^, with minor modifications outlined below. Briefly, full length SMCHD1 and Lamin B1 sequences were PCR amplified from oligo-dT-primed cDNA and cloned into pENTR-vector (Thermo Fisher Scientific). The SMCHD1 and Lamin B1 cDNAs were subsequently cloned into pLgw EcoDam-V5-RFC1 ^65^ (a gift from Bas van Steensel, Addgene plasmid # 59209), to construct pLgw EcoDam-V5-SMCHD1 and EcoDam-V5-Lamin B1 by gateway cloning. The expression of EcoDam-V5-SMCHD1 fusion protein (Dam-SMCHD1) and EcoDam-V5-Lamin B1 fusion protein (Dam-LaminB1) were confirmed by Western blot in LHCN-M2 cells transiently transfected with pLgw-EcoDam-V5-SMCHD1 or pLgw-EcoDam-V5-Lamin B1. To generate stable cell lines with low expression of Dam, Dam-SMCHD1 and Dam-Lamin B1, respectively, the strong CMV promoter was removed by using the Q5 Site-Directed Mutagenesis Kit (NEB, E0554S), to ensure that the fragments harboring the Dam fusion protein were driven by the HSP70 promoter. The puromycin resistance gene was then cloned into the vector for screening sExtended Data Table ingle clones. Stably transfected cells were selected in 1 µg/mL puromycin (Thermo Fisher Scientific, A1113803) for 10 days, then single clones of the transfected cell were selected and maintained with 1 µg/mL puromycin. Genomic DNA of stable lines was purified using Quick-DNA Miniprep Plus kit (Zymo Research, D4070) with the RNase A digestion step included. The RNase treated genomic DNAs were then digested with DpnI overnight (at least 12 h) at 37°C. We cleaned up the DpnI-digested DNA with the Qiagen PCR Purification Kit according to the manufacturer’s instructions. We ligated the DamID adaptors to 750 ng (up to 15 µl) DpnI-digested DNA for 2 h at 16°C (15 μL DpnI-digested DNA, 2 μL 10×T4 DNA ligase buffer, 0.8 μL dsAdR, 1.2 μL H_2_O), followed by 10 min incubation at 65 °C to inactivate the ligase. We added 19 μL of premade TaDa DpnII digestion buffer (4 μL 10× DpnII buffer, 15 μL H_2_O) and 1 μL of DpnII enzyme into the mixture and digested the mixture at 37°C for 3 hours. We added 118 μL premade DamID PCR buffer (16 μL 10×cDNA buffer, 2.5 μL DamID_PCR primer (50 μM), 3.2 μL 10 mM dNTPs, 96.3 μL water) and 2 μL of Advantage2 cDNA polymerase enzyme to the DpnII-digested DNA, split the PCR reaction into 4×40 μL reactions, then perform PCR with the programs as followed: 68°C for 10 min, 1 cycle of 94°C for 30 sec, 65°C for 5 min, 68°C for 15 min. Then we added 3 cycles of 94°C for 30 sec, 65°C for 1 min, 68°C for 10 min, and 17 cycles of 94°C for 30 sec, 65°C for 1 min, 68°C for 2 min, followed by a final extension step with 68°C for 5 mins. We purified PCR products with Qiagen PCR Purification Kit according to the manufacturer’s instructions. We sonicated 2 μL of purified DNA with a Covaris E220 Evo sonicator to shear the DNA to 300 to 500 bp size fragments. We incubated 1 μL of AlwI enzyme (NEB, R0513S) and sheared mixtures at 37°C for overnight to remove the DamID adaptors. We cleaned up the mixtures with AMPure Beads according to the manufacturer’s instructions. Then, we incubated 500 ng of cleaned DNA in 20 μL with 2.5 μL end-repair enzyme mix (1.14 μL T4 DNA polymerase (3 U/μL), 0.23 μL Klenow fragment (5 U/μl), 1.14 μL T4 polynucleotide kinase) and 7.5 μL of end-repair buffer (3 μL 10× T4 DNA ligase buffer, 1.2 μL 10 mM dNTPs, 3.3 μL H_2_O) at 30 °C for 30 mins. Adenylation of 3ʹ ends was then performed by incubating the mixture with 0.75 μl of Klenow 3ʹ– 5ʹ exo^-^ enzyme at 37°C for 30 min followed by incubating with 2.5 μL of sequencing adaptor and 2.5 μL of NEB quick ligase enzyme at 30 °C for 30 min. Two rounds of AMPure beads cleanup were then performed according to the manufacturer’s instructions. The purified DNA was then amplified by PCR using the following conditions: 98°C for 30 sec, 6 cycles of 98°C for 10 sec, 60°C for 30 sec, 72°C for 30 sec, then a final extension at 72°C for 5 mins. Sequencing was performed with an Illumina HiSeq 2500 instrument with 150-bp paired-end read runs.

#### In *situ* Hi-C library preparation and sequencing

In situ Hi-C was performed as described previously ^42,66^. Briefly, 5 × 10^6^ cells were cross-linked with 1% formaldehyde for exactly 10 min at room temperature. We added 1.25 mL of 2.5 M glycine to quench the cross-linking reaction for 5 min at room temperature, and then placed the cells on ice for at least 15 min. We pelleted the cells by centrifugation at 1000×g for 10 min. We washed the pellet with DPBS (Thermo Fisher Scientific,14190-144), then cross-linked with disuccinimidyl glutarate (Thermo Fisher Scientific, 20593, final concentration 3 mM) for 40 min at room temperature. Cross-linked cells were lysed in ice-cold lysis buffer (10 mM Tris-Cl, pH 8.0, 10 mM NaCl, 0.2% Igepal CA-630, MP Biomedicals, 198596) containing 10 μL protease inhibitor cocktail. The nuclei were digested with 400 U DdeI and 400 U DpnII (NEB) at 37°C overnight with interval shaking (900 rpm, 30 sec on, 4 min off). We incubated the Hi-C samples with biotinylation buffer (7 μL 10x NEBuffer 3.1, 1.5 μL 10 mM dCTP, 1.5 μL 10 mM dGTP, 1.5 μL 10 mM dTTP, 37.5 μL 0.4 mM biotin-14-dATP, 11 μL H_2_O, 10 µL Klenow DNA polymerase) for 4 hours at 23°C in a thermomixer (900 rpm, 30 sec on, 4 min off). We incubated the ligation mix (240 μL 5x ligation buffer, 120 μL 10% Triton X-100, 12 μL 10 mg/ml BSA, 243 μL H_2_O, 50 μL 1 U/ μL T4 DNA ligase) with the biotinylation mix for 4 hours at 16°C in a thermomixer with interval shaking (900 rpm, 30 sec on, 4 min off). To each Hi-C sample, we added 50 μL of 10 mg/ml proteinase K and incubated for 2 hours at 65°C with interval shaking (900 rpm, 30 sec on, 4 min off), then added another 50 μL of 10 mg/ml proteinase K to each Hi-C sample and continued incubating overnight at 65°C. We prepared biotin removal reactions in PCR tubes as follows: 15 μg of Hi-C DNA sample, 13 μL 10x NEB buffer 2.1, 3.25 μL 1 mM dATP, 3.25 μL 1 mM dGTP, 13 μL 1 mM 3000 U/ml T4 DNA polymerase and filled up to 130 μL with water. We transferred two 65 μL aliquots from each 130 μl reaction to a thermocycler with the following cycling parameters: 20°C for 4 hours, 75°C for 20 min, hold at 4°C. We sonicated the samples on a Covaris E220 Evo sonicator to shear the DNA to a fragment size distribution of 300 to 500 bp. We performed double size selection on the sheared DNA using 0.8X-1.1X Agencourt AMPure XP beads (Beckman Coulter, A63881) according to the manufacturer’s instructions. We incubated end-repair mix (7 μL 10× ligation buffer, 7 μL 2.5 mM dNTP mix, 2.5 μL 3 U/μL T4 DNA polymerase, 2.5 μL 10 U/μl T4 polynucleotide kinase, 0.5 μL 5 U/μl Klenow DNA polymerase, 4.5 μL H_2_O) and size-selected HiC libraries in a thermocycler with the following cycling parameters: 20°C for 30 min, 75°C for 20 min, hold at 4°C. The products were then purified with MyOne Streptavidin C1 beads (Thermo Fisher Scientific, 65001), dA-tailed, and ligated to 5 μL 15 μM adaptor at 37°C for 2 hours. Hi-C libraries were then amplified by PCR using the following conditions: 98°C for 30 sec, 4 cycles of 98°C for 10 sec, 65°C for 30 sec, 72°C for 30 sec, then a final extension at 72°C for 2 mins. In situ Hi-C on WT and SMCHD1 knockout LHCN-M2 cells were performed in two biological replicates, which were sequenced on an Illumina Novaseq 6000 machine with 150 bp paired-end read runs.

#### Electron microscopy

Our present study employed a comprehensive methodology to prepare and visualize cellular ultrastructure using transmission electron microscopy (TEM). To ensure precise cellular preservation, cells were initially fixed using a solution composed of 2% paraformaldehyde and 2.5% glutaraldehyde in a 0.1 M cacodylate buffer at room temperature for 2 hours. Following the fixation step, the cells underwent a series of washing steps with 0.1 M cacodylate buffer (3 washes of 20 minutes each) to remove any residual fixative. Subsequently, post-fixation was carried out by immersing the cells in a solution containing 1% osmium tetroxide and 1.25% potassium ferrocyanide in a 0.1 M cacodylate buffer for 1 hour at room temperature. After several buffer washes, the cells were immersed in a mixture of 50% ethanol and 1% uranyl acetate for 30 minutes as part of the en-bloc staining process. Dehydration was then performed by exposing the cells to ascending ethanol concentrations (50%, 70%, 90%, and 95%), ultimately reaching 100% absolute ethanol. The cells were treated with propylene oxide for 20 minutes. Infiltration of the cells was achieved by immersing them in a mixture of LX-112 resin (Ladd Research, USA, 21210) and propylene oxide, with varying ratios (1:3, 1:1, 3:1) for 2 hours each and overnight in 100% resin. Samples were placed in a 60°C oven for polymerization for 3 days. Ultrathin sections (∼ 70 nm) were obtained using a Leica Ultracut UCT ultramicrotome and mounted on copper-rhodium grids. Post-staining was carried out with aqueous uranyl acetate and Reynolds’ lead citrate. The high-resolution imaging of the prepared samples was conducted using a Tecnai Spirit G2 BioTWIN transmission electron microscope operating at 120 kV, facilitated by a digital capture system utilizing an Orius 832 CCD Camera and Gatan software.

### Quantification and Statistical Analysis

#### RNA-seq data analysis

For RNA-seq analysis, we followed our standard procedures ^30,67^. Seventy-five base pair reads were trimmed with Trim Galore (v0.4.0), trimmed reads were then aligned to the human genome hg19 with STAR (version 2.5.1), and gene count was performed with STAR. Differential gene expression was called by the Limma (v3.38.2) statistical package19. P-values for differential expression were adjusted for multiple testing correction using the Benjamini-Hochberg method in the stats package. Statistical significance for differentially expressed genes was fold change >2 with q <0.05. Pheatmap package was used to generate heat maps. The GO function annotation was performed with PANTHER (v18.0). The motif analysis for the promoter of the differentially expressed genes in this study was performed with ‘findMotifsGenome.pl’ function of Homer (v4.11.1).

#### WGBS data analysis

For WGBS data analysis, following our standard procedures ^30,67^, 150 bp paired-end whole-genome bisulfite reads were trimmed using TrimGalore (v0.5.0) with the following parameters to remove library preparation artifacts and low quality bases: –-length 50, –-clip_R1 10, –-clip_R2 18, –-three_prime_clip_R1 10, and –-three_prime_clip_R2 10. Trimmed reads were aligned to the reference genome (hg19) using Bismark ^68^ (v0.19.0) and Bowtie2 ^69^ (v2.3.3.1) with the following parameters: –X 1000, –-nucleotide_coverage, and –-bowtie2. Duplicates were marked and removed using the deduplicate_bismark script provided with Bismark. CpG methylation values were extracted using the bismark_methylation_extractor script provided with Bismark and the following parameters: –-no_overlap, –-comprehensive, –-merge_non_CpG, and –-cytosine_report. We used DMRseq ^70^ version 0.99.0 to identify DMRs. Briefly, CpG loci with fewer than three reads were not considered for DMR calling, and a single CpG coefficient cutoff of 0.05 was used for candidate regions. Significant DMRs were identified using a q value < 0.05. Genomic coordinates of Hypermethylation-DMR clusters (Hyper-DMR clusters) or Hypomethylation-DMR clusters (Hypo-DMR clusters) were then defined based on Hyper-DMRs (Hypo-DMRs) called from the last step. Consecutive Hyper-(or Hypo-) DMRs smaller than 10 kb bins were merged. At least five merged Hyper-DMRs (or Hypo-DMRs) were considered as a Hyper-DMR cluster (or Hypo-DMR cluster) for further analysis. These DMR clusters are listed in Extended Data Table 2.

#### ChIPseq data analysis

The adapter and low-quality sequences were trimmed from 3ʹ and 5ʹ ends by Trimgalore (v0.5.0). Subsequently, the trimmed reads were aligned to hg19 with Bowtie2 (versionv2.5.1). The aligned reads were then deduplicated using PicardTools (http://broadinstitute.github.io/picard). ChIP-peaks were then analyzed with MACS2 ^71^. All ChIP data was normalized against the corresponding input controls using the ‘-c’ option of MACS2. For CTCF, RAD21, H3K4me3 and H3K27ac, ChIP-Seq peaks were called using the ‘callpeak’ function of MACS2 with default parameters. For H3K9me3, H3K36me2, H3K36me3, H3K4me1 and Pol II, the ‘–broad’ option of MACS2 was used to identify broad peaks. Bigwig tracks of ChIP-seq were generated using bamCoverage of deeptools ^72^ (v3.5.5) to compute coverage across the entire genome. Heatmaps and enrichment profiles of average Counts Per Million mapped reads (CPM) were performed using deeptools. Overlap of genomic regions and peaks contained within them was determined with the bedtools (v2.30.0). Enrichment patterns of different ChIPseq data was generated by genomation ^73^ (v1.34.0) and deeptools.

#### Hi-C data analysis

The in situ Hi-C data were processed with a standard pipeline of Juicer ^74^, using a pipeline code available at (https://github.com/theaidenlab). Within sample normalization was performed using the Knight-Ruiz (KR) method ^75^. We generated Hi-C data comprised of two sets of biological replicates for each WT and SMCHD1 knockouts. Each map contains an average of 2.4 billion contacts with mapping quality (MAPQ) score >30. The contacts correlation analysis for biological replicates was performed at 500 kb resolution with R package multiHiCcompare (v 1.20.0) ^76^, then the two biological replicates for each genotype were merged to get higher resolution (average of 4.8 billion contacts) for further analysis. For each chromosome in each sample, compartments were called using the ‘eigenvector’ option of Juicer tools (https://github.com/theaidenlab). The eigenvector was calculated to delineate compartments, which was the first principal component of the Pearson’s correlation matrix of the Observed/Expected, and the sign of the first eigenvector was used to assign compartment labels. We used H3K9me3 and H3K27ac data to flip the sign of the eigenvector such that the value of the first eigenvector was larger than 0 indicating an active A compartment and values smaller than 0 indicating an inactive B compartment. The heatmap and line plot of the AB correlation matrix eigenvector was generated by FAN-C ^77^ (v 0.9.1) based on the PC1 information generated by Juicer.

HiCCUPS (https://github.com/theaidenlab) was used to identify the chromatin loops with default parameters for high resolution maps in HiCCUPS described with all the available flags (-m 512 –c (all chromosomes) –r 5000,10000 –k KR –f.1,.1 –p 4,2 –i 7,5, –t 0.02,1.5,1.75,2 –d 20000,20000,50000), as described previously ^78^. All the code used in the above steps is publicly available at (https://github.com/theaidenlab). Juicebox ^74^ was used for visualization of the Hi-C data. All the code used in the above steps is publicly available at (https://github.com/theaidenlab).

To show the average contact count distribution of loops and their surroundings between WT and SMCHD1 knockouts, we performed an aggregate peak analysis (APA) of loop calls using the R package GENOVA ^79^ (v1.0.0.9.98) at 10 kb resolution. To show the average contact count distribution of differential TADs and their surroundings in WT and SMCHD1 knockouts, we performed an aggregated TAD analysis (ATA) using the ATA-tool from the R package GENOVA (v1.0.0.9.98), including the up and downstream 500 kb regions, from Hi-C contact maps at 10 kb resolution.

#### Analysis of SMCHD1 DamID-seq datasets

DamID-seq 150 bp paired end reads were trimmed with Trim Galore (v0.5.0). The trimmed reads were then aligned to hg19 with Bowtie2 (v2.5.1). To get the normalization of signals generated by Dam-fusion proteins over background, the aligned reads were binned into GATC fragments according to GATC sites, then were normalized to Dam-only control, using damidseq_pipeline ^80^. The log2 (Dam-fusion/Dam alone) values of each GATC fragment was then calculated. To identify all Dam-SMCHD1 and Dam-Lamin B1 binding sites, all Dam-fusion data was normalized against the corresponding Dam-only control using the ‘-c’ option of MACS2 with major parameters – broad, –min-length 50000, –-max-gap 50000 and –broad-cutoff 0.0001.

#### Electron microscopy analysis

Quantifications of TEM images were performed using Fiji version (2.9.0) by thresholding the 32-bit TEM images. The outer border of each nucleus was traced using the freehand tool, here a stylus pen on a touch screen for precise selection. The heterochromatin was then identified by invert and binary functions, and the measurements > threshold limit was set to measure the area of the heterochromatin near the nuclear periphery and the total nuclear threshold. The percentage of lamina-adjacent heterochromatin was calculated by dividing the lamina-adjacent heterochromatin by the total nuclear area. Finally, the relationship between normal and FSHD2 samples, as well as between wildtype and SMCHD1-depleted LHCN-M2 cells was plotted based on the percentage of heterochromatin. Independent data from three samples with N = 30, 20, and 20 nuclei for normal myoblasts and N = 20, 20, and 20 nuclei for FSHD2 myoblasts were obtained, as well as three different clones with N = 19, 20, and 20 nuclei for LHCN-M2 WT cells and N = 19, 19, and 20 nuclei for the SMCHD1 mutant LHCN-M2 cells. Visualization of the data and statistical testing were performed using R package (version 4.1.2).

#### 3D genome modeling

3D genome models for WT and SMCHD1-KO LHCN-M2 cells were generated with Chrom3D (v1.0.2) ^43^, following a protocol described previously, with minor modifications ^44^. Briefly, the TADs were generated using the Arrowhead algorithm ^78^, which was used in Chrom3D simulations as chains of contiguous beads for modeling chromosomes. Centromeres and assembly gaps were filtered from the TAD data which were subsequently used for the next step. LADs (LaminB1-DamID data generated in this study) were matched to corresponding TADs and used as peripheral sub-nuclear constraints in the simulations. The significant intra-chromosomal and inter-chromosomal interactions were identified with the non-central hypergeometric distribution algorithm, part of Chrom3D. Gtrack files then were generated from significantly interacting TADs and LADs using the Python scripts makeGtrack.py, which were used as inputs for the next 3D simulation step with Chrom3D. Chrom3D simulations were performed for each condition with a nuclear occupancy of 0.05, with 2 million simulation steps and parameter –-nucleus, which was set to push the beads towards the nuclear interior. The nuclear boundary was modeled as a sphere with a radius of 5 μm. The simulated Chrom3D models then were visualized with ChimeraX^81^

**Extended Data Fig. 1:**
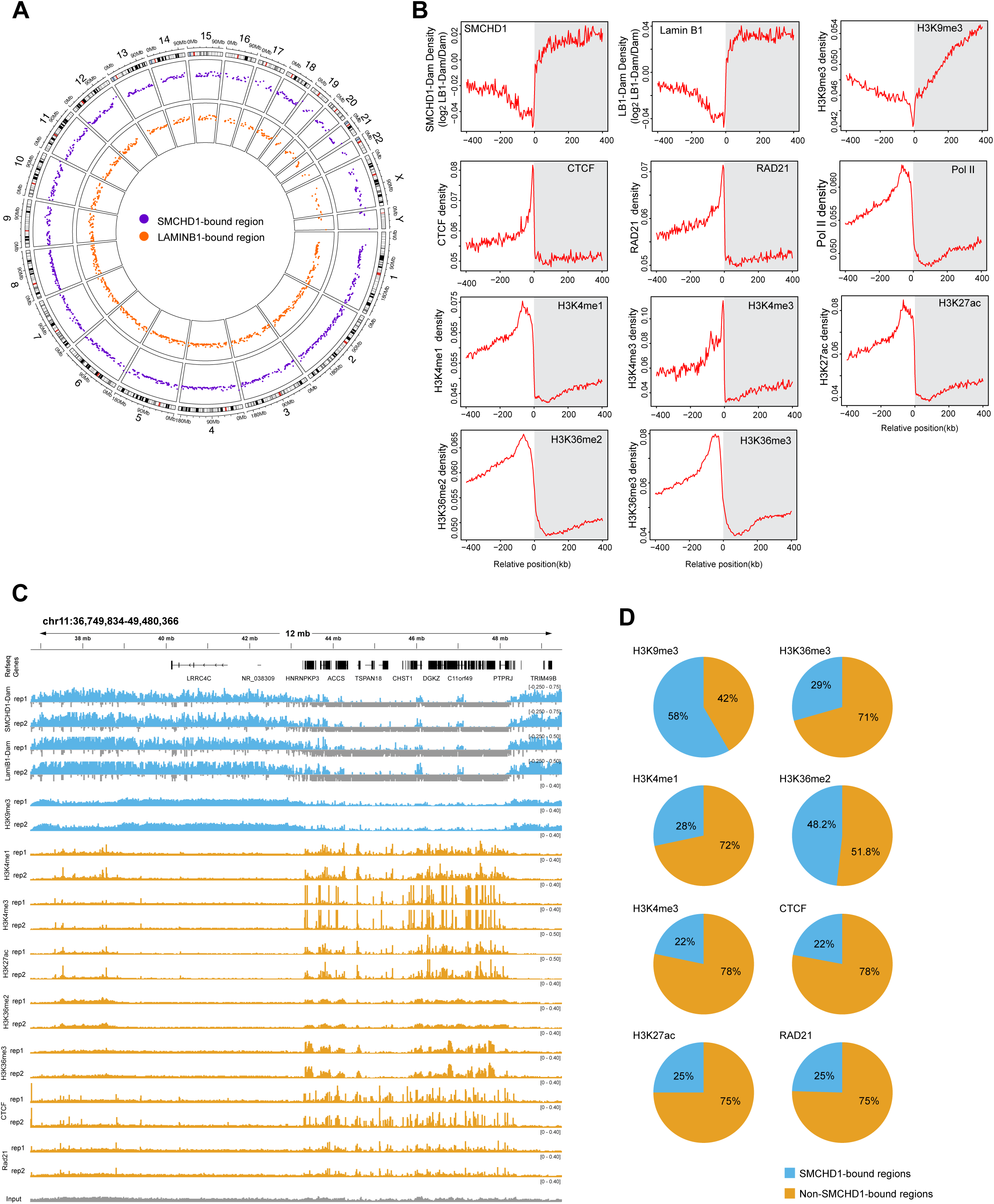
Relationship between SMCHD1 and LADs. **A**. Genome-wide colocalization of SMCHD1 (blue) and Lamin B1 (red) as shown by Circos plot. **B.** Transition of active and inactive (SMCHD1, Lamin B1, H3K9me3) marks at LAD borders. The active marks are depleted within the LADs (grey shaded areas) and the inactive components are enriched. **C.** Mapping of SMCHD1, Lamin B1 and several histone modifications along an area of chromosome 11. This segment of the chromosome contains a region bound by SMCHD1 (left side) and a region that is not bound (right side). **D.** Global distribution of different histone modifications over regions bound by SMCHD1 (blue) or not bound by SMCHD1 (orange).

**Extended Data Fig. 2:**
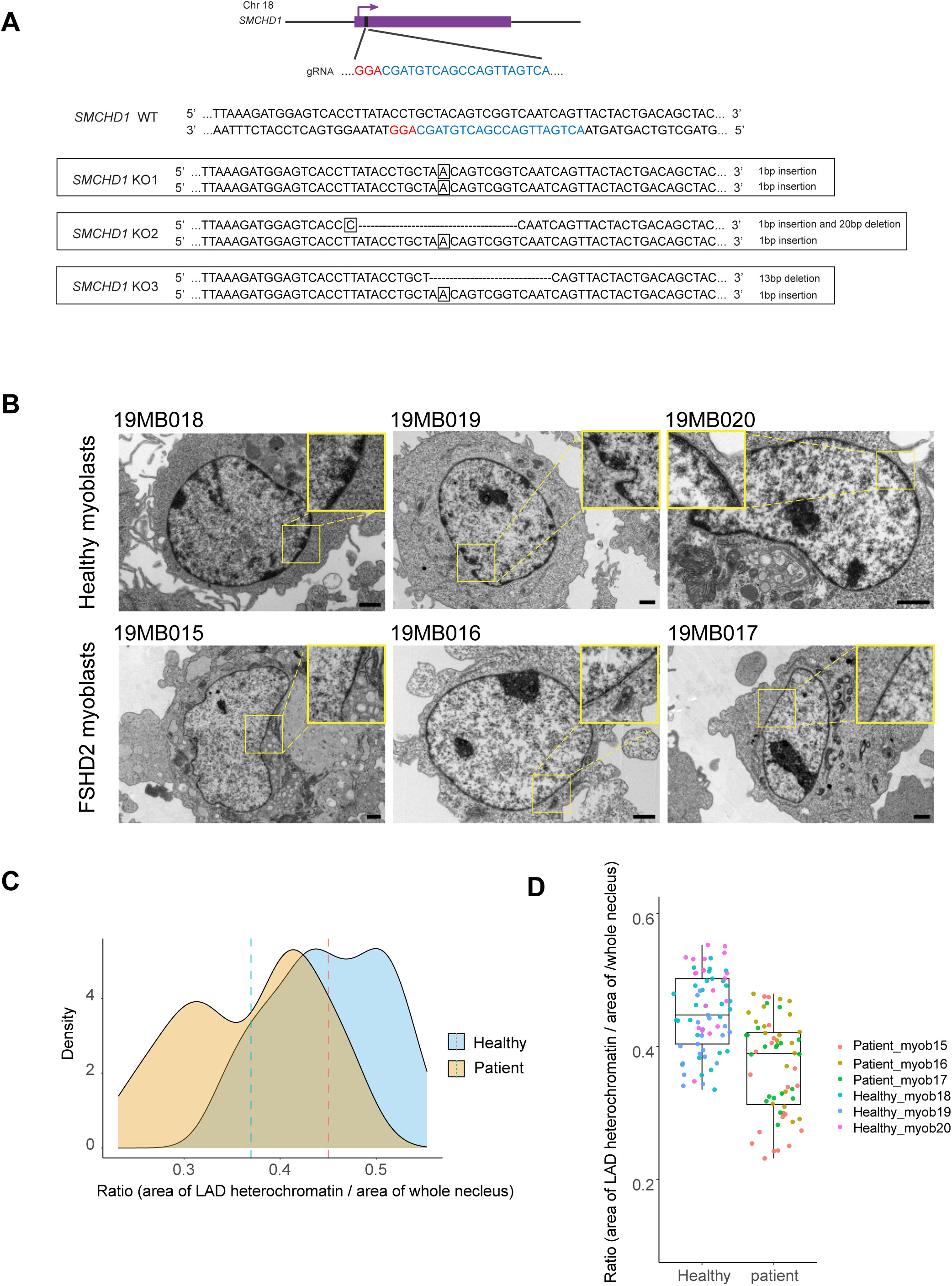
Genotypes of SMCHD1 knockouts and transmission electron microscopy (TEM) of nuclear lamina-associated heterochromatin in healthy myoblasts and myoblasts from SMCHD1-deficient FSHD2 patients. **A**. Genotyping of three CRISPR/Cas9 knockout clones used in this study. The small guide RNA targeting region is in the third exon of human *SMCHD1*. Sanger sequencing confirmed the presence of biallelic frame-shift mutations. Sequences targeted by the guide RNA are colored in blue, and the PAM sequence is colored in red. **B.** TEM images of myoblasts from healthy individuals and from patients with FSHD2. The insets show a magnification of specific areas of the nuclear periphery. **C, D.** Quantitation of perinuclear heterochromatin from TEM images in healthy and FSHD2 cells. Regions under the nuclear envelope were quantitated by densitometry of 70 healthy and 60 FSHD2 cells. The heterochromatin staining intensities were normalized to the total area of the nucleus and show a reduction in FSHD2 cells (P = 2.184e-10, t-test).

**Extended Data Fig. 3:**
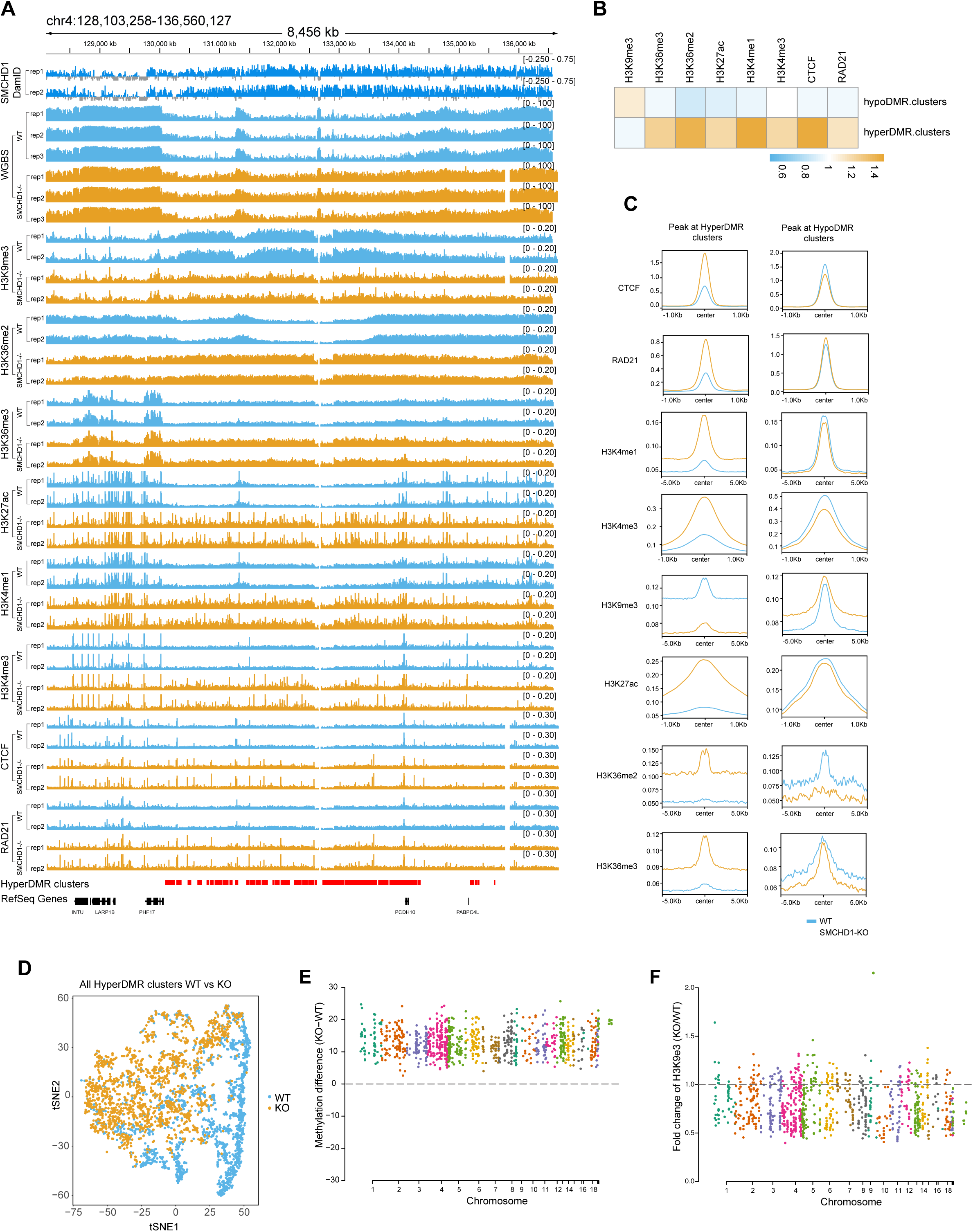
Extensive changes of heterochromatin and euchromatin marks and structural proteins at clustered DNA hypermethylated regions. **A**. Genome browser tracks of SMCHD1, DNA methylation (WGBS), H3K9me3, H3K36me2, H3K36me3, H3K27 acetylation, H3K4me1, H3K4me3, CTCF and the cohesin subunit RAD21 are displayed. Two tracks are shown for each mapped parameter. Hypermethylation cluster regions and Refseq genes are shown at the bottom. **B.** The heatmap shows the global differences for inactive and active chromatin marks, CTCF, RAD21, and RNA polymerase II over all clusters of DNA hypermethylation and DNA hypomethylation. **C.** Composite profile plots of the indicated histone modifications and structural proteins over the center of DNA hypermethylation clusters (left) and DNA hypomethylation clusters (right). Blue lines, wildtype myoblasts; orange lines, SMCHD1^−/−^ myoblasts. **D.** tSNE plots of HyperDMR clusters at the whole genome level in WT and SMCHD1^−/−^ cells. All HyperDMR clusters were plotted based on combinatorial mean levels of H3K9me3, H3K4me1, H3K4me3, H3K27ac, H3K36me2, H3K36me3, CTCF, RAD21 and RNA polymerase II across the clusters in two dimensions, using t-Distributed Stochastic Neighbor Embedding (t-SNE). **E.** Differences in methylation (%) at hypermethylation DMR clusters in SMCHD1-bound regions between KO and WT cells for all chromosomes. Each dot represents a hypermethylation DMR cluster. **F.** Differences in H3K9me3 at the same hypermethylation DMR clusters as shown in panel E.

**Extended Data Fig. 4:**
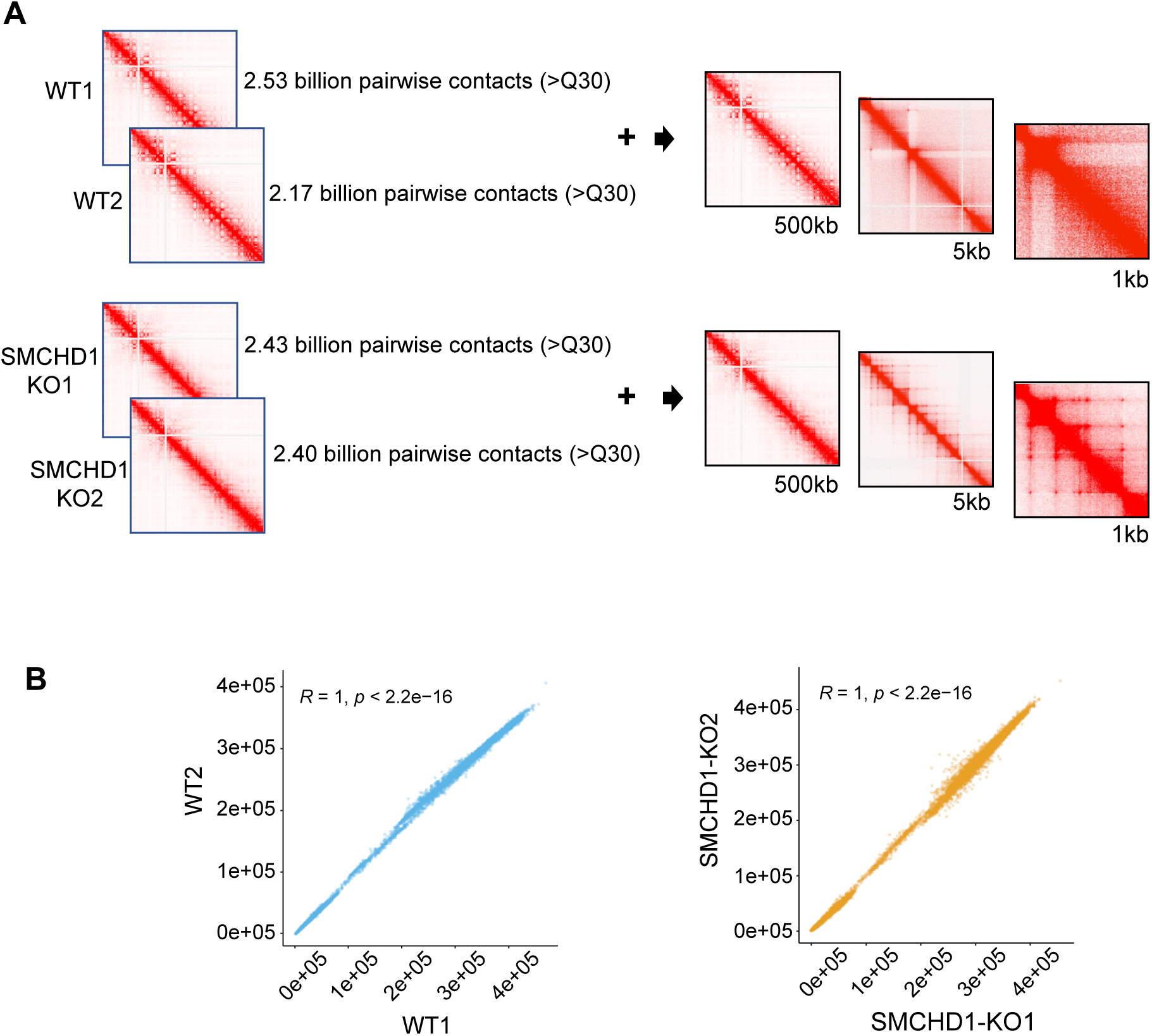
Generation of HiC 3.0 data. **A**. Representation and resolution achieved in interaction maps. **B.** Correlation between biological replicates. Global HiC contacts were calculated at 500 kb resolution.

**Extended Data Fig. 5:**
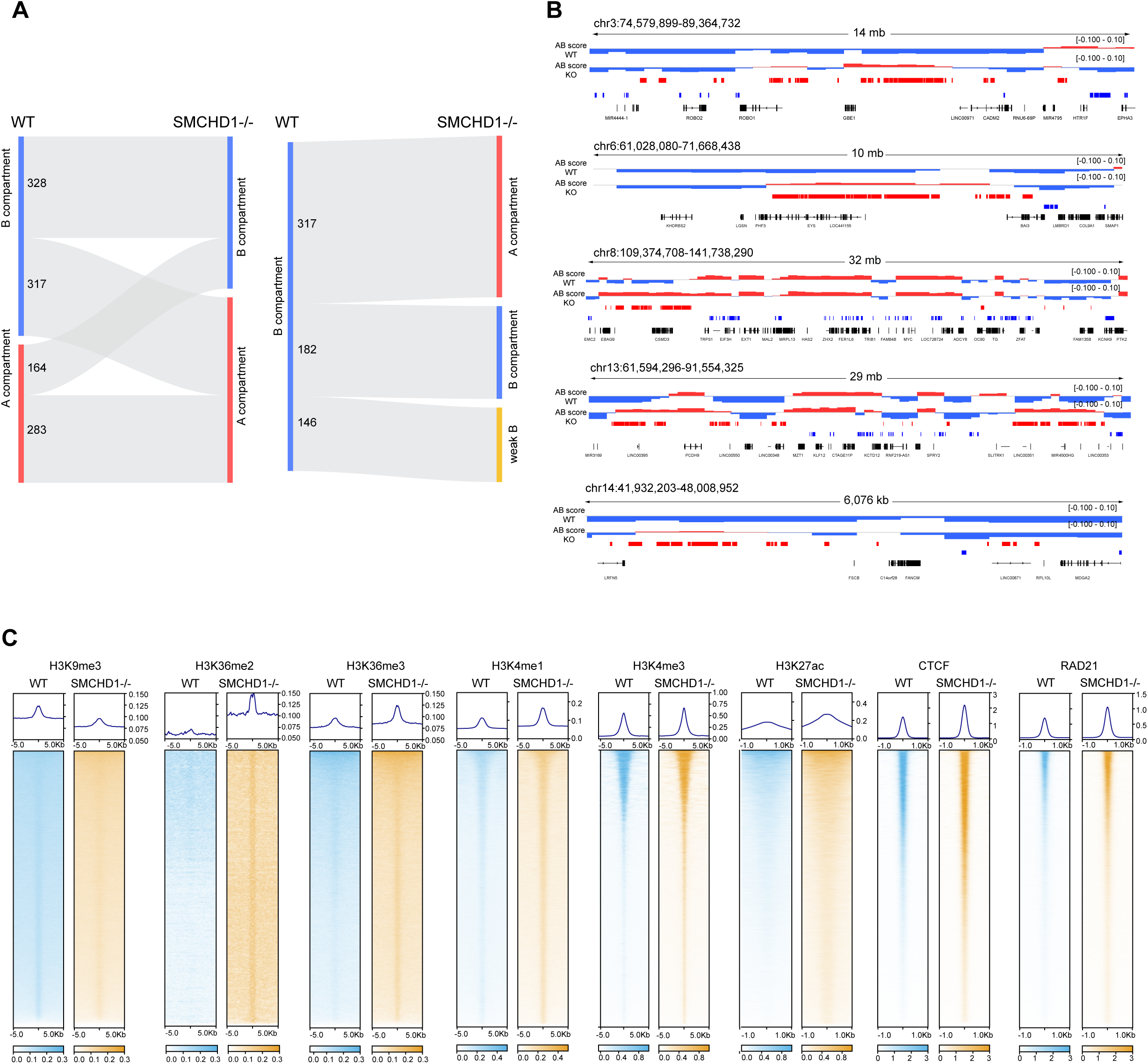
Changes in genome compartmentalization after loss of SMCHD1. **A**. Sankey Plot showing the number of A and B compartment transitions after SMCHD1 inactivation (left panel). The number of B compartments transitioned to weak B compartment are shown in the right panel. **B.** Examples of B-to-A transitioned regions on four different chromosomes. Several B compartments (blue) are changing to A compartments (red) or weak B compartments (thin blue lines). **C.** Composite profile plots and heatmaps showing histone modifications, CTCF and RAD21 over all B-to-A transitioned regions.

**Extended Data Fig. 6:**
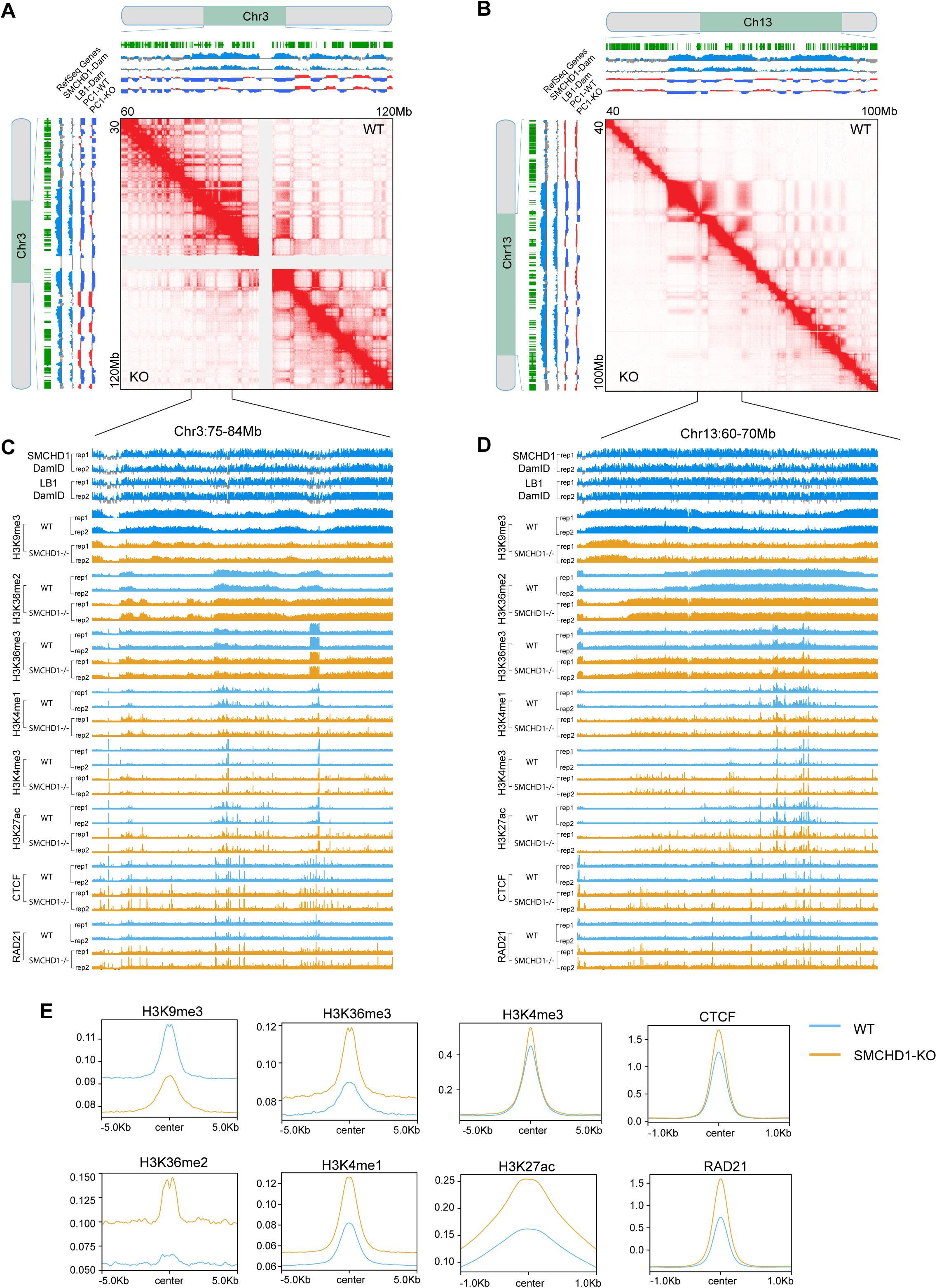
Loss of contacts between B compartments in SMCHD1-depleted cells and histone modification patterns over regions of B compartment loss. **A**. In situ Hi-C contact map showing contacts between individual B compartments (red stripes of dots, blue to blue contacts in the PC1 maps) across a 60 Mb region of chromosome 3 in WT and SMCHD1^−/−^ cells. Many of these contacts are not found in the absence of SMCHD1. The SMCHD1 and Lamin B1 signals are shown along with B compartments (blue) and A compartments (red) on the top and left. **B.** In situ Hi-C contact map showing contacts between individual B compartments (blue to blue in the PC1 maps) across a 60 Mb region of chromosome 13 and their loss in SMCHD1^−/−^ cells. The SMCHD1 and Lamin B1 signals are shown along with B compartments (blue) and A compartments (red) on the top and left. **C.** SMCHD1, Lamin B1, CTCF, RAD21, and histone modification patterns in WT and SMCHD1^−/−^ cells in a magnified representative region from panel A. **D.** SMCHD1, Lamin B1, CTCF, RAD21, and histone modification patterns in WT and SMCHD1^−/−^ cells in a magnified representative region from panel B. **E.** Plots showing the differential histone mark enrichment patterns across all B compartment loss regions at the whole genome scale.

**Extended Data Fig. 7:**
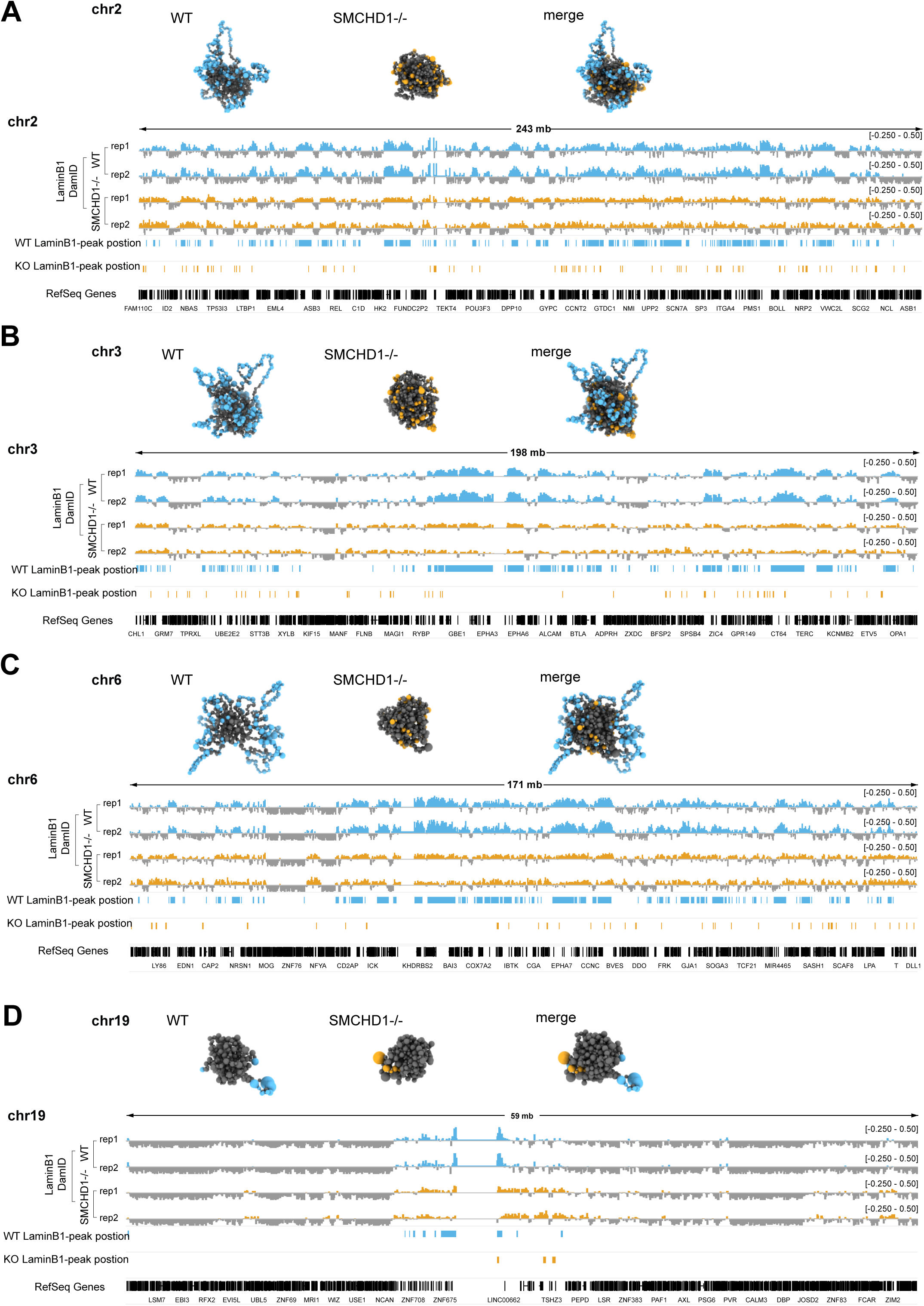
Chrom3D analysis of single chromosomes in WT and SMCHD1^−/−^ cells. **A**. Chromosome 2. The Lamin B1 sequence tracks, peak positions and Refseq genes are shown at the bottom. Light blue, Lamin B1 associated regions in wildtype cells; orange, Lamin B1 regions in SMCHD1 knockout cells. **B.** Chromosome 3. **C.** Chromosome 6. **D.** Chromosome 19, a gene-rich chromosome with few LADs.

**Extended Data Fig. 8:**
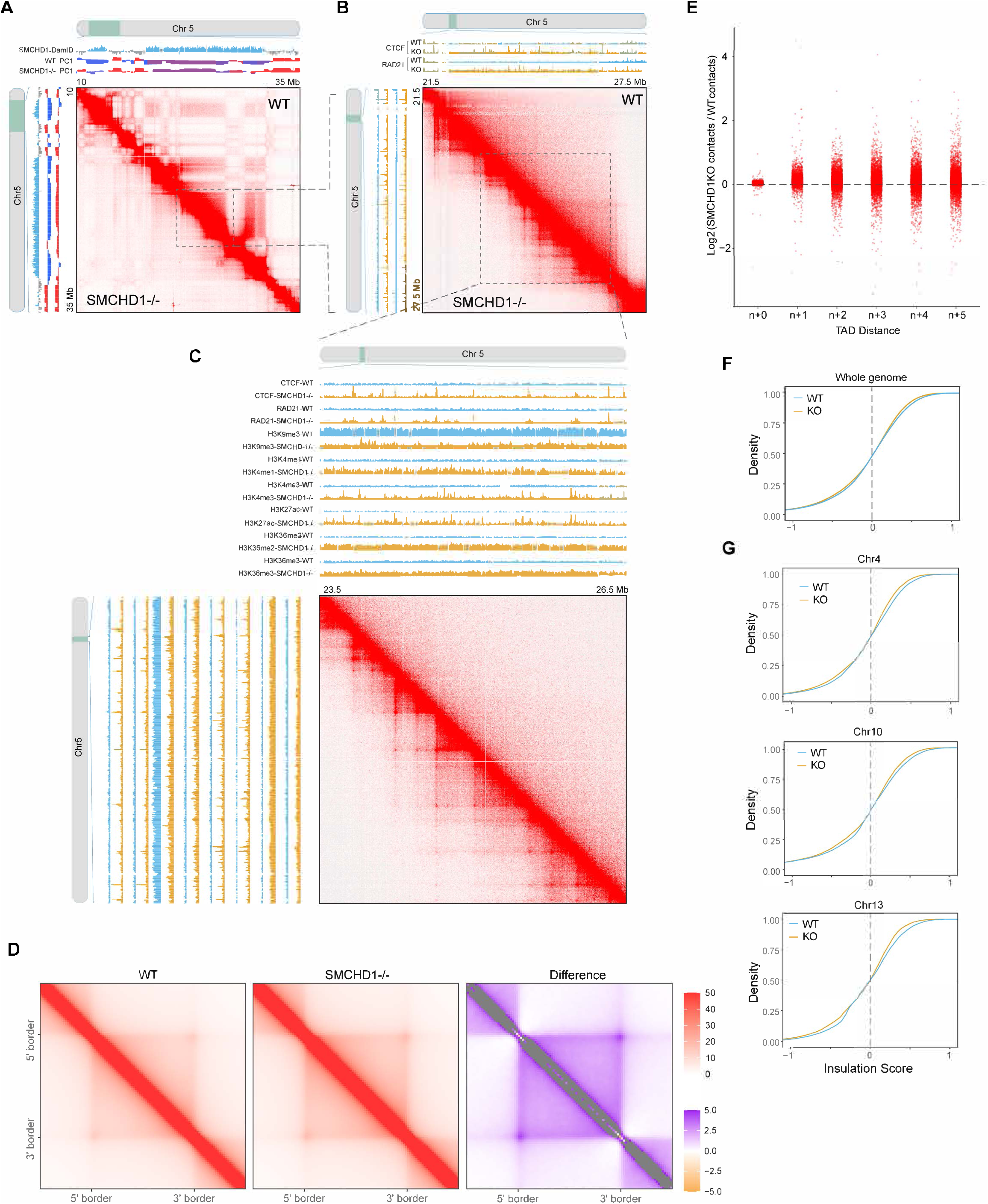
Gains of TADs and loops in SMCHD1-deficient cells. **A**. Hi-C contact maps showing contacts along a presentative region on chromosomes 5 in wildtype (WT) and SMCHD1^−/−^ cells. The SMCHD1-bound regions and compartments are shown alongside the heatmap. **B.** Zoomed-in view of the interactions shown in panel A. Regions are examples of genomic locations harboring increased TADs in SMCHD1^−/−^ cells. Differential ChIP-seq patterns of CTCF and RAD21 are shown at the top and left of the heatmap. **C.** Zoomed-in view and browser tracks of interactions shown in panel B. Regions are examples of genomic sites harboring increased loop domains at new TADs in SMCHD1^−/−^ cells. Differential binding patterns of CTCF and RAD21, and histone marks are shown at the top and left of the heatmap. **D.** Aggregate TAD analysis (ATA) of TADs flanked by CTCF binding sites detected in wildtype and SMCHD1^−/−^ cells shows increased interaction intensity in SMCHD1^−/−^ cells versus wildtype cells for B-to-A compartment transitions. A differential ATA analysis plot with purple denoting more interactions in SMCHD1^−/−^ cells compared to wildtype cells is shown on right. **E.** TAD+N analysis computes the interaction density within TADs and their 1, 2,., N neighbors. This analysis can be used to compare whether TADs in two samples interact differently with their neighboring TADs. **F.** Cumulative distribution plots of the insulation scores at the whole genome in wildtype and SMCHD1^−/−^ cells as indicated (P<2.2e-16, Kolmogorov-Smirnov test). **G.** Cumulative distribution plots of the insulation scores on presentative chromosomes 4, 10 and 13 in WT and SMCHD1^−/−^ cells as indicated (P<2.2e-16 for chr4, P=1.797e-08 for chr10, P=8.993e-15 for chr13, Kolmogorov-Smirnov test).

**Extended Data Fig. 9:**
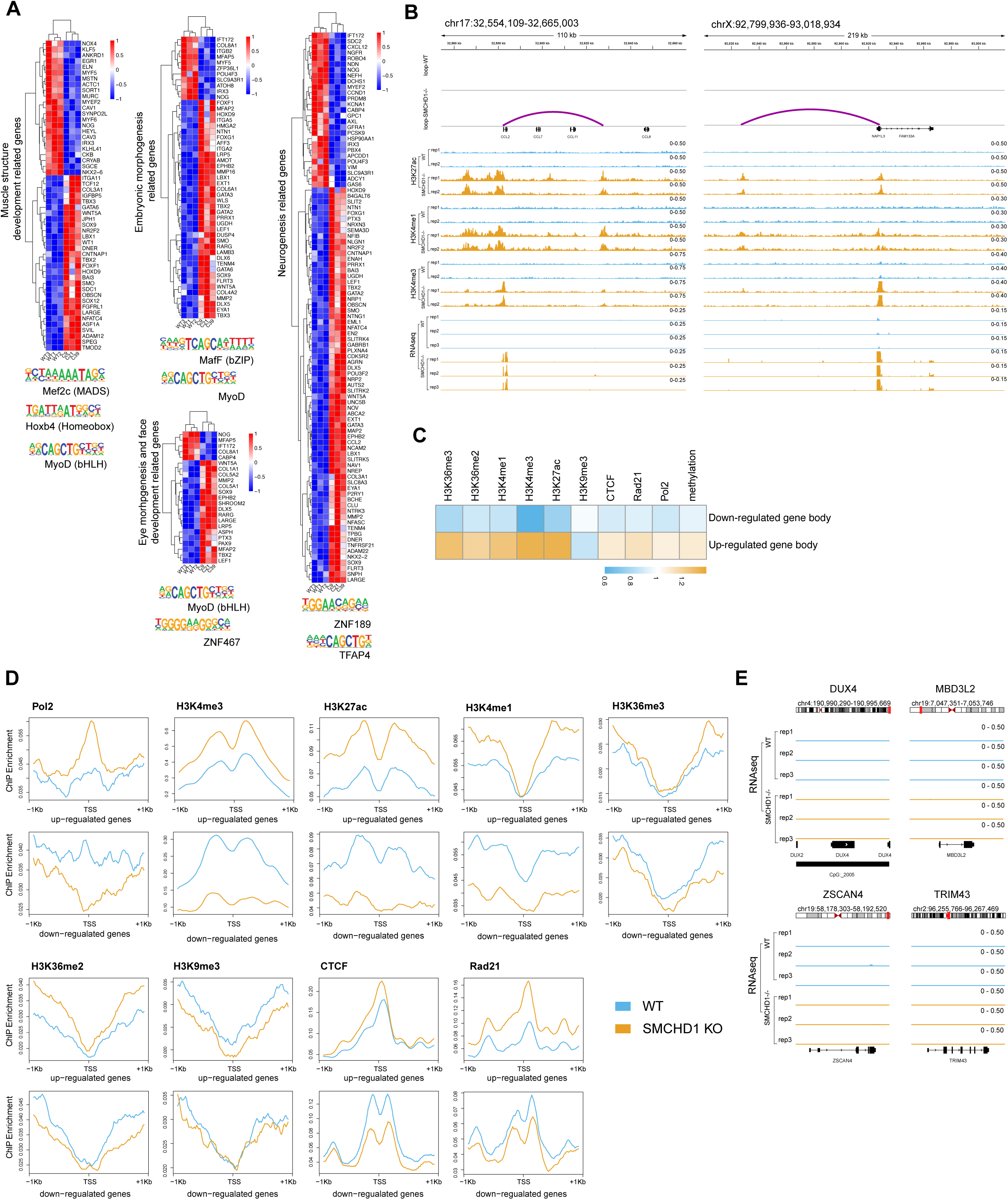
Gene expression changes after inactivation of SMCHD1. **A**. Heatmaps of upregulated and downregulated genes after inactivation of SMCHD1. The genes are grouped according to gene ontology terms. Putative transcription factor motifs found in the promoters (TSS+/− 1kb) of these genes are shown at the bottom of the heatmaps. **B.** Examples of gene activation events linked to formation of new enhancer-promoter loops and gain of active enhancer and promoter chromatin marks. *CCL2*, left panel. *NAP1L3*, right panel. **C.** Relationship of upregulated or downregulated genes with gain or loss of chromatin marks along the gene body (TSS to TES). **D.** Levels of histone modifications and DNA binding proteins over the TSS of all differentially expressed genes (including upregulated genes and downregulated genes) in wildtype and SMCHD1-deficient cells. DEGs were called with fold change >2, FDR <0.05. **E.** Expression of *DUX4* and several DUX4 target genes in wildtype and SMCHD1-mutant myoblasts as analyzed by RNA-seq analysis.

**Extended Data Fig. 10:**
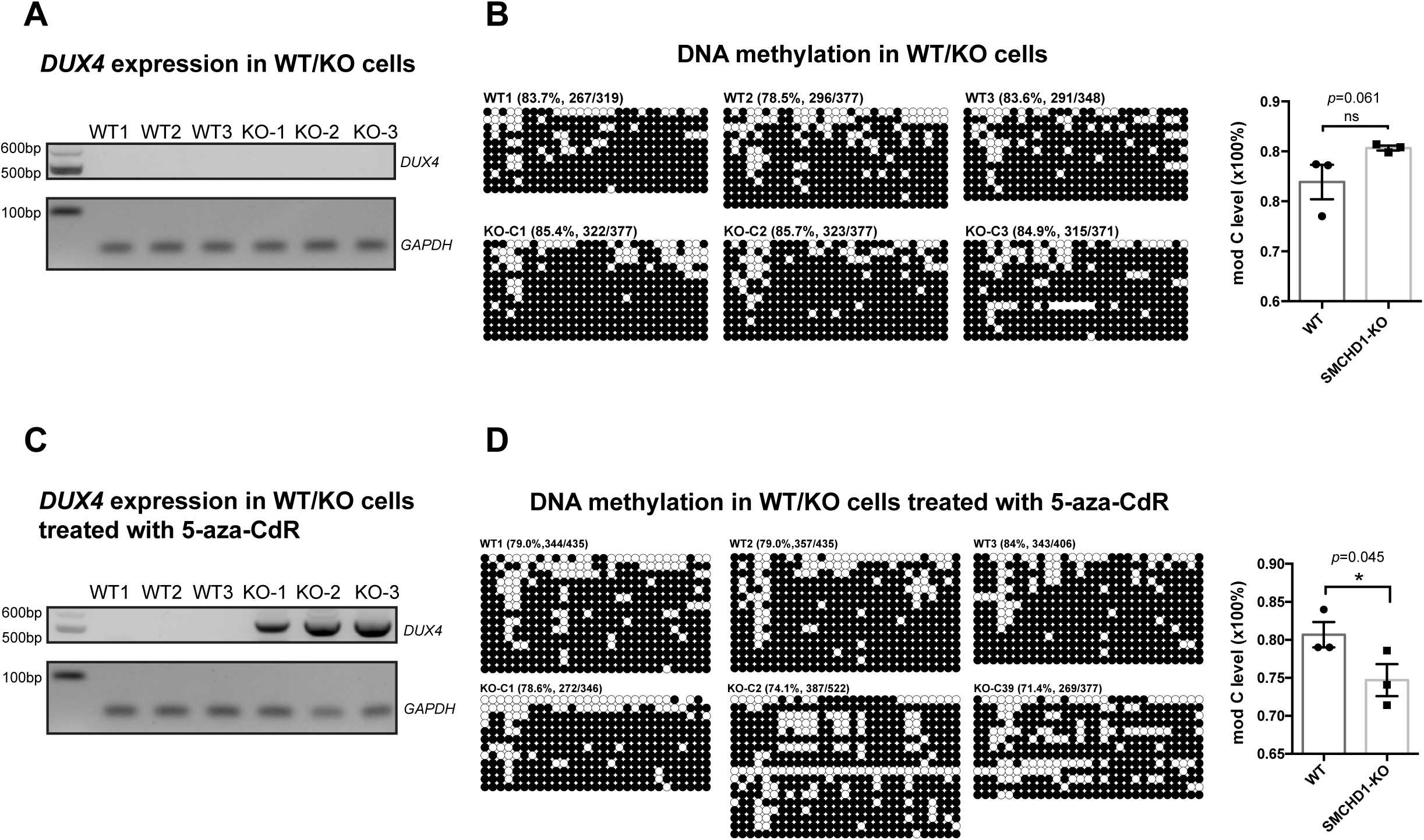
Expression and DNA methylation changes of *DUX4* in myoblasts after inactivation of SMCHD1. **A**. RT-PCR of *DUX4* expression in wildtype (WT) and SMCHD1 knockout (KO) cells. **B.** DNA methylation at the *DUX4* locus 1 kb upstream of the TSS as determined by manual bisulfite sequencing. Black circles represent methylated CpG sequences, open circles are unmethylated CpGs. The percentage of modified CpGs is indicated. **C.** RT-PCR of *DUX4* expression in wildtype (WT) and SMCHD1 knockout (KO) cells treated with 5-zazC-dR. **D.** DNA methylation at the *DUX4* locus after 5-azaC-dR treatment as determined by manual bisulfite sequencing. Black circles represent methylated CpG sequences, open circles are unmethylated CpGs. The percentage of modified CpGs is indicated.

## Supplementary Movies

**Movie S1: Chromosome configuration in WT myoblasts**.

Radius is 5 μm.

**Movie S2: Chromosome configuration in SMCHD1-KO myoblasts**.

Radius is 5 μm.

**Movie S3: Spherical view of chromosome and LAD configuration in WT myoblasts**.

The LAD-associated genomic regions are indicated as blue spheres. Grey spheres represent TADs without LAD association. Radius is 5 μm.

**Movie S4: Spherical view of chromosome and LAD configuration in SMCHD1-KO myoblasts.**

The LAD-associated genomic regions are indicated as blue spheres. Grey spheres represent TADs without LAD association. Radius is 5 μm.

**Movie S5: Cross-sectional view of chromosome and LAD configuration in WT myoblasts**.

The LAD-associated genomic regions are indicated as blue spheres. Grey spheres represent TADs without LAD association. Radius is 5 μm.

**Movie S6: Cross-sectional view of chromosome and LAD configuration in in SMCHD1-KO myoblasts**.

The LAD-associated genomic regions are indicated as blue spheres. Grey spheres represent TADs without LAD association. Radius is 5 μm.

## Notes

### Competing Interest Statement

The authors have declared no competing interest.

